# Exercise remodels hippocampal extracellular matrix to alleviate chondroitin-4-sulphate–induced memory impairment

**DOI:** 10.1101/2025.11.14.688432

**Authors:** Natalie E. Doody, Ben Stevens, Nicole J. Smith, Varinder K. Lall, Sally Boxall, Ruth Hughes, Elizabeth C. Akam, Graham N. Askew, Jessica C. F. Kwok, Ronaldo M. Ichiyama

## Abstract

Research utilising exercise therapeutically to enhance neuroplasticity mainly reports enhanced expression of neurotrophic (neural growth) factors which are associated with increased neurogenesis, structural changes, and improved memory.

However, these neural changes are tightly regulated by both promotors and inhibitors of neuroplasticity, and the effect of exercise on inhibitory pathways is currently understudied.

We show that exercise also modulates inhibitors of neuroplasticity. Six weeks of treadmill training reduced the gene expression of aggrecan, a chondroitin sulphate proteoglycan (CSPG), and altered the composition of CSPG containing structures, perineuronal nets, in the rodent hippocampus. Genetically manipulating chondroitin-4-sulphation of CSPGs by overexpressing hippocampal *Chst11* impaired memory performance, which was mitigated by treadmill training.

We provide evidence that hippocampal CSPGs are involved in object recognition memory, and that there is a link between exercise, modulating hippocampal inhibitors of neuroplasticity, and memory performance.

## Introduction

Exercise is a potent lifestyle intervention that improves memory and enhances neuroplasticity, such as neurogenesis, dendritic spine density and long term potentiation (Fernandes et al., 2020, Hötting and Röder, 2013), which is important in healthy ageing and neurorehabilitation. It is widely accepted that exercise exerts its structural and functional neuroplastic effects via the upregulation of the neurotrophic factor, brain derived neurotrophic factor (BDNF). However, the mechanisms of exercise-induced plasticity and memory improvements remain elusive. Whilst BDNF levels are fundamental in promoting neuroplasticity, there are inhibitory modulators of neuroplasticity in the adult central nervous system which work to maintain optimum plasticity and normal physiological functions (Oberman and Pascual-Leone, 2013). To focus on BDNF alone omits a significant part of the plasticity regulating mechanism, and it is possible that exercise simultaneously upregulates neurotrophic factors and downregulates inhibitory modulators of plasticity to create a more permissive and flexible neural environment. Fully elucidating the exercised-induced regulation of neuroplasticity will be advantageous for improving neuroprotection, neurorehabilitation, and cognition, and expanding the therapeutic potential of exercise.

Inhibitory modulators of plasticity refer to molecules which restrict cytoskeletal/anatomical arrangements (e.g. growth cone collapse, neurite outgrowth, reorganisation of synapses, and dendritic spine formation) (Lai and Ip, 2013, Baldwin and Giger, 2015, Boghdadi et al., 2017, Sorg et al., 2016, Akbik et al., 2012). This study refers to three subtypes of inhibitory modulators of neuroplasticity: chondroitin sulphate proteoglycans (CSPGs), myelin-associated inhibitors, and negative guidance molecules. These inhibitory modulators of neuroplasticity are extracellular stimuli which converge to activate the RhoA/ROCK signalling pathway, which is known to induce growth cone collapse, restrict neurite outgrowth, and negatively regulate dendritic spine formation (Spence and Soderling, 2015, Fujita and Yamashita, 2014, Doody et al., 2024). In rodent models, exercise training has been shown to downregulate the myelin associated inhibitors, Nogo-A and MAG, in the hippocampus after traumatic brain injury, and reduce Nogo-A in the cortex and striatum following ischaemic stroke (Gomez-Pinilla et al., 2008, Zhang et al., 2013). In the healthy central nervous system, research on exercise-induced regulation of inhibitory modulators of plasticity is extremely limited, however, voluntary wheel running reduced the level of CSPG containing structures, perineuronal nets (PNNs), in the hippocampus (Smith et al., 2015, Terstege et al., 2024).

PNNs are extracellular structures which surround the cell soma and proximal dendrites of neuronal subtypes (predominantly parvalbumin interneurons) and are known to restrict neuroplasticity in development, central nervous system injury, and in memory systems (Li et al., 2024). CSPGs are main components of PNNs and consist of core proteins to which chondroitin sulphate glycosaminoglycan (CS-GAG) chains attach. The degradation of CS-GAGs has been documented to improve object recognition memory, learning acquisition, and fear extinction (Gogolla et al., 2009, Romberg et al., 2013, Carulli et al., 2020). Furthermore, the sulphation pattern of CS-GAGs determines how they affect neuroplasticity. Chondroitin-4-sulphation (C4S) (which is upregulated by *Chst11*) is known to be inhibitory, whilst chondroitin-6-sulphation (C6S) (which is upregulated by *Chst3*) is more permissive and plasticity promoting (Wang et al., 2008, Lin et al., 2011).

In the present study, we aimed to determine whether exercise training had the capacity to downregulate inhibitory modulators of plasticity in the healthy central nervous system and whether this was associated with improved memory. We trained Wistar rats in a moderate intensity treadmill training paradigm (30 minutes/day, 5 days/week, for six weeks); an exercise paradigm which is more translatable to human health than voluntary wheel running. Our data shows that treadmill training downregulated gene expression of the CSPG encoding gene *Acan*, altered the biochemical composition of hippocampal PNNs by reducing CS-GAG levels, and attenuated memory deficits which were induced by virally overexpressing *Chst11* in the hippocampus. This highlights a novel mechanistic narrative for how exercise training enhances memory function. CS-GAGs act as molecular brakes on neuroplasticity which can be lifted by an exercise regime that can easily be prescribed to humans for improving memory and supporting healthy aging. Furthermore, CS-GAGs are potential therapeutic targets for exercise mimetics and the pharmaceutical treatment of cognitive decline.

## Results

### Exercise training downregulated RhoA/ROCK pathway components in the hippocampus, specifically, Acan

To investigate whether exercise had the capacity to modulate inhibitors of neuroplasticity throughout the central nervous system, we trained seven rats to run on a motorised treadmill in a six-week exercise programme, whilst seven rats remained sedentary. As previous research in this area is limited, we applied an exploratory, high through-put approach using RT-qPCR to determine whether 25 genes within the RhoA/ROCK pathway were susceptible to changes following exercise training. The RhoA/ROCK pathway restricts neuroplasticity by regulating dendritic spine morphology, actin cytoskeleton dynamics, and induces growth cone collapse thus inhibiting neurite outgrowth (Fujita and Yamashita, 2014, Spence and Soderling, 2015).

Exercise training downregulated the overall gene expression profile of inhibitory RhoA/ROCK pathway components in the hippocampus, whilst the cortex and lumbar region of the spinal cord remained stable (Figure 1A). Exercise training significantly downregulated the relative mRNA expression of hippocampal *Acan* in comparison to the sedentary cohort (Exercise: - 0.6078 ± 0.17 Log2 FC compared to sedentary control; t_(4)_=2.880, *p*=0.045) (Figure 1B). *Acan* encodes aggrecan, one of the CSPGs, which are known to bind to two receptors within the RhoA/ROCK pathway, NgR1 and NgR3 (Dickendesher et al., 2012). Full statistical analyses for gene expression data in the hippocampus, cortex, and lumbar regions are presented in Extended Data Table 1.

**Figure 1.**
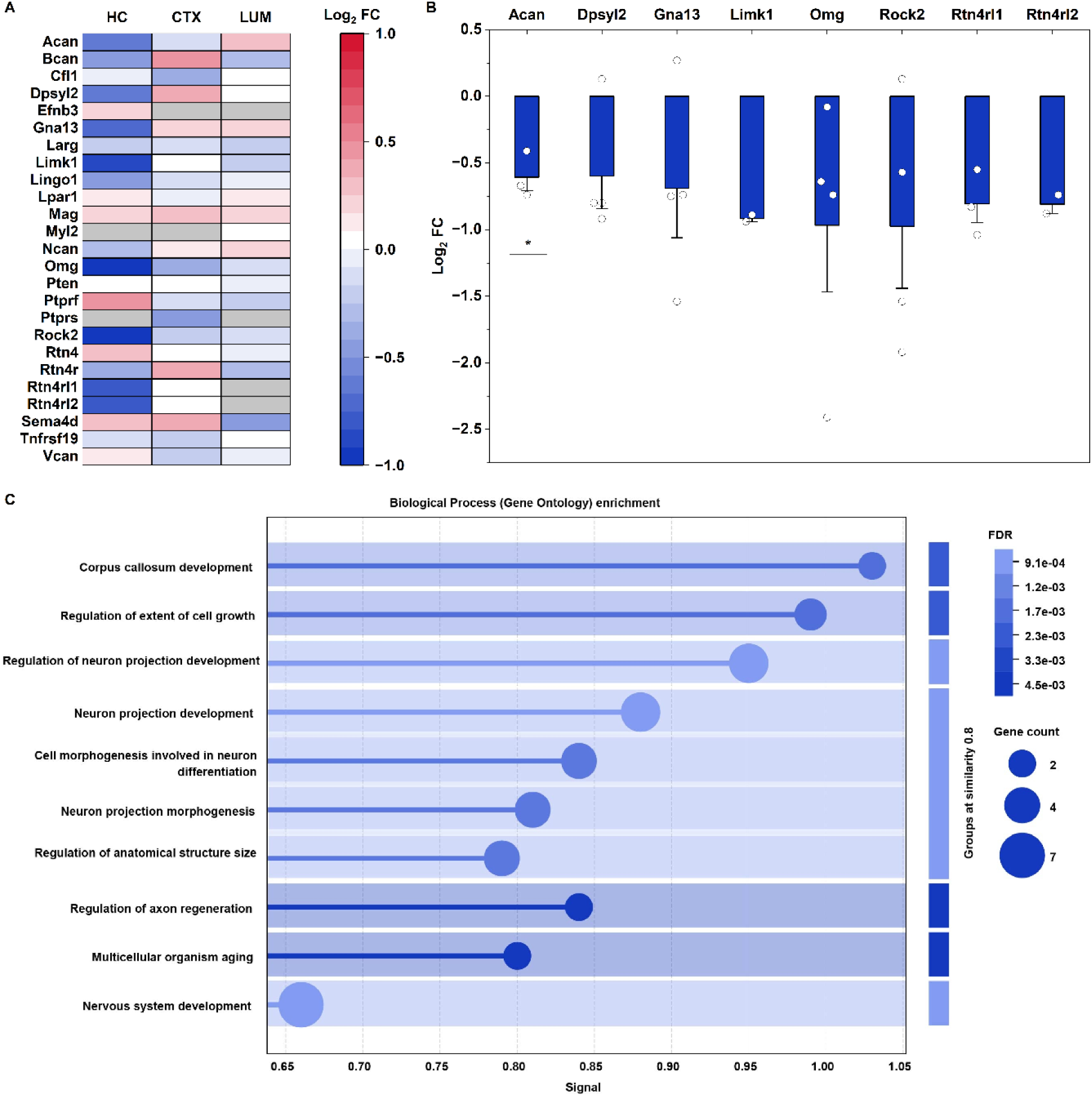
Exercise training downregulates RhoA/ROCK pathway genes in the hippocampus. **A.** Heatmap presenting mean relative fold change (Log2 FC) of RhoA/ROCK pathway gene expression following exercise training. HC – hippocampus; CTX – cortex; LUM – lumbar. Blue - downregulated mRNA expression, red - upregulated mRNA expression, grey - no remaining data point after exclusion criteria applied. **B.** Hippocampal genes with Log2 FC values between the cut off points of ≤0.75 and ≥1.5 following exercise training. Statistical significance determined using an independent t test. * *p*<0.05 when compared to sedentary control. Exercise training significantly downregulated *Acan* mRNA (*p*=0.045). Bars represent mean relative fold change of mRNA expression compared to sedentary controls. Error bars: SE. *n*=3/4 except *Limk1* and *Rtn4rl2* with *n*=2 due to exclusion criteria. **C.** STRING database enrichment analysis: Top 10 Gene Ontology terms associated with the gene list from B. Size of circle represents number of genes aligned with the Gene Ontology term. FDR – false discovery rate (p value corrected by Benjamini-Hochberg procedure).

Although only one statistically significant gene change was identified in the hippocampus, all target genes are components of the same biological pathway, and thus several smaller gene changes may cumulatively impact overall RhoA/ROCK signalling. Hence, we applied relative fold change cut off values of ≤0.75 and ≥1.5 to the RT-qPCR data to investigate the gene expression profiles more thoroughly. After these cut off points were introduced, exercise training downregulated eight genes including: two inhibitory ligands which can activate the RhoA/ROCK pathway, *Acan* and *Omg;* two Nogo receptors, *Rtn4rl1* and *Rtn4rl2;* the effector molecule *Rock2*; and the downstream signalling molecules *Gna13, Limk1,* and *Dpysl2* (Figure 1B).

We used the online STRING database to perform enrichment analysis of the protein-protein interactions of the eight identified genes to gain further functional insight into these exercise-modulated inhibitory networks. The enrichment analysis showed a significant number of interactions between the proteins corresponding to our exercise-regulated gene list (protein-protein interaction enrichment: *p* = 0.00000382). The enrichment analysis aligned the gene list to two neuroplasticity related KEGG Pathways: ‘Axon guidance’ (rno04360) and ‘Regulation of actin cytoskeleton’ (rno04810). The top 10 Gene Ontology Biological Processes are presented in (Figure 1C).

In summary, we have shown that the inhibitory RhoA/ROCK pathway, known to restrict neuroplasticity, is susceptible to modulation by exercise training, and have identified *Acan* as a novel candidate in exercise-induced neuroplasticity.

### Exercise training modified the composition of hippocampal perineuronal nets

We next sought to characterise morphological changes related to the candidate *Acan* in the hippocampus following exercise training. *Acan* encodes the CSPG, aggrecan, which is found in the loose extracellular matrix and is also a main component of PNNs (Rowlands et al., 2018). PNNs are lattice-like structures which mainly surround parvalbumin interneurons, preventing synaptic reorganisation, and thus reducing neuroplasticity (Sorg et al., 2016). Structurally, aggrecan consists of a core protein to which CS-GAG side chains are covalently bonded (Figure 2A). We used an anti-aggrecan antibody to label the aggrecan core protein and *Wisteria floribunda* agglutinin (WFA) to label CS-GAG chains. This allowed us to visualise the two different components of CSPGs, and classify whether hippocampal PNNs were one of three different phenotypes: 1) PNNs containing both aggrecan core protein and CS-GAG chains (ACAN+/WFA+); 2) PNNs containing only aggrecan core protein (ACAN+/WFA-); and 3) PNNs containing CS-GAG chains but not aggrecan core protein (ACAN-/WFA+).

**Figure 2.**
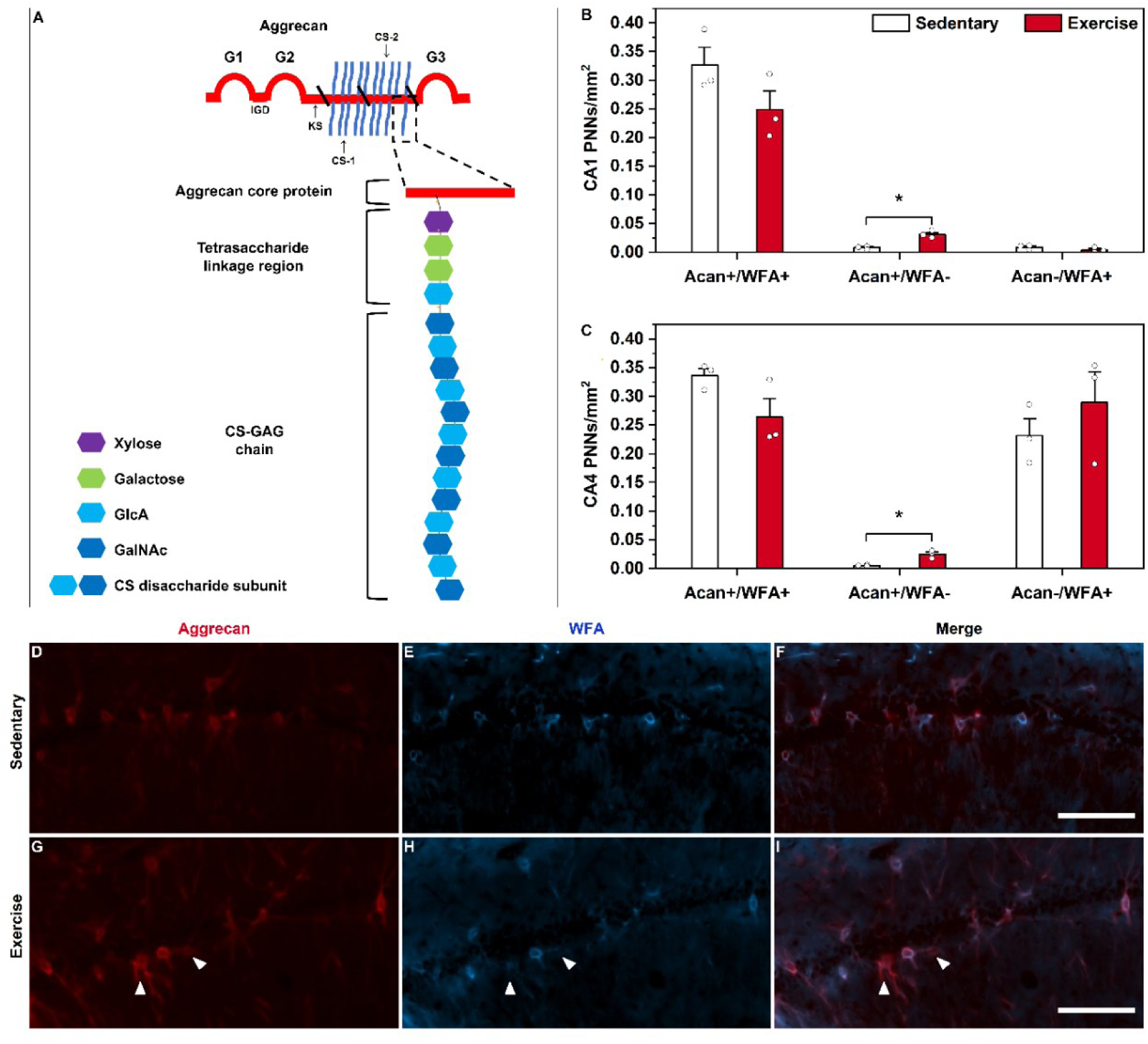
Exercise training reduced the level of CS-GAGs in hippocampal PNNs. **A**. Schematic of the chondroitin sulphate proteoglycan (CSPG), aggrecan. CSPGs consist of a core protein (depicted in red) to which chondroitin sulphate glycosaminoglycan (CS-GAG) chains (depicted in blue) attach to. CS-GAG chains covalently attach to serine residues in the core protein by a tetrasaccharide linkage region containing xylose, two galactose residues, and glucuronic acid (GlcA). The chondroitin sulphate chain is then extended with repeating disaccharide subunits of GlcA and *N*-acetylgalactosamine (GalNAc). G1-3: globular domains. IGD: interglobular domain. KS: keratan sulphate attachment domain. CS-1 and CS-2: chondroitin sulphate attachment domains **B-C**. Perineuronal nets (PNNs) were counted as one of three different phenotypes: labelled with aggrecan and WFA (ACAN+ WFA+), labelled with aggrecan only (ACAN+ WFA-), or labelled with WFA only (ACAN-WFA+).ACAN: aggrecan antibody which labelled the core protein. WFA: *Wisteria floribunda* agglutinin which labelled CS-GAG chains. Statistical significance was assessed using independent t tests. * *p*<0.05 when compared to sedentary control. Exercise training increased the number of PNNs which contained only aggrecan core protein without CS-GAG chains (ACAN+/WFA-) in CA1 (*p*=0.005) and CA4 regions (*p*=0.009). Bars represent mean PNN count. Error bars: SE. *n*=3. **D-I**. PNNs labelled with aggrecan (**D** and **G**), WFA (**E** and **H**), and a MERGE of PNNs labelled with both aggrecan and WFA (**F** and **I**) in CA1 region of sedentary and exercised animals. White arrowheads indicate PNNs that express aggrecan without WFA following exercise training. Scale bars: 50 μm.

The total number of PNNs, calculated as the sum of all three phenotypes, was comparable between sedentary and exercise trained animals in all hippocampal subfields (CA1, CA2, CA3, CA4, dentate gyrus (DG)) (Extended Data Table 2). However, exercise training altered the morphological composition of PNNs. Exercise training significantly increased the number of PNNs that contained only aggrecan core protein without CS-GAG chains (ACAN+/WFA-) in CA1 (Sedentary: 0.0076 ± 0.0018 PNNs/mm^2^; Exercise: 0.0310 ± 0.0037 PNNs/mm^2^; t_(4)_=-5.674, *p*=0.005) and CA4 (Sedentary: 0.0052 ± 0.0006 PNNs/mm^2^; Exercise: 0.0250 ± 0.0041 PNNs/mm^2^; t_(4)_=-4.778, *p*=0.009) (Figure 2B-C). The total number of PNNs in these hippocampal subfields did not change, suggesting that exercise training reduced the level of CS-GAG chains within aggrecan-positive PNNs in CA1 and CA4. PNN phenotype counts for CA2, CA3, and DG subfields are presented in (Extended Data Figure 1). Whilst exercise training altered the composition of hippocampal PNNs, PNN thickness (both aggrecan and WFA) remained unchanged between trained and sedentary animals (Extended Data Figure 2).

As CSPGs, such as aggrecan, are also expressed in the loose extracellular matrix we measured the staining intensity of anti-aggrecan and WFA in the hippocampus. In all hippocampal subfields, the staining intensity of aggrecan and WFA was comparable between exercised and sedentary animals (Extended Data Figure 3).

This is the first study to show that exercise in the form of treadmill training modulates the biochemical composition of PNNs, known plasticity regulators, in the hippocampus. Without changing the total number of PNNs, exercise training induced a shift towards a higher proportion of PNNs containing only aggrecan and lacking WFA labelling. This suggests that exercise training downregulated CS-GAGs specifically in aggrecan-positive PNNs, potentially increasing hippocampal neuroplasticity.

### Exercise training mitigates memory impairment induced by chondroitin-4-sulphation

The enzymatic degradation of CS-GAGs (which cleaves CS-GAGs from CSPG core proteins such as aggrecan) has been shown to improve the acquisition of new memories (Romberg et al., 2013, Carulli et al., 2020). As we have demonstrated that exercise training modulated CS-GAGs in hippocampal PNNs, we next sought to determine whether this was associated with improved memory. The removal of PNNs and CS-GAGs are common approaches to assess their function in memory processes, however, we required a contrasting experimental paradigm to mechanistically test whether the exercise-induced modulation of CS-GAGs improved memory. CS-GAG chains contain repeated disaccharide subunits of glucuronic acid and N-acetylgalactosamine. These disaccharide subunits can be sulphated at several different positions which affects how they regulate neuroplasticity. Chondroitin-4-sulphate (C4S) contains sulphation at the 4^th^ position of N-acetylgalactosamine within the disaccharide unit (Figure 3C). C4S is known to be restrictive to neuroplasticity and memory, whereas its counterpart chondroitin-6-sulphate (C6S) is plasticity promoting (Wang et al., 2008, Lin et al., 2011).

**Figure 3.**
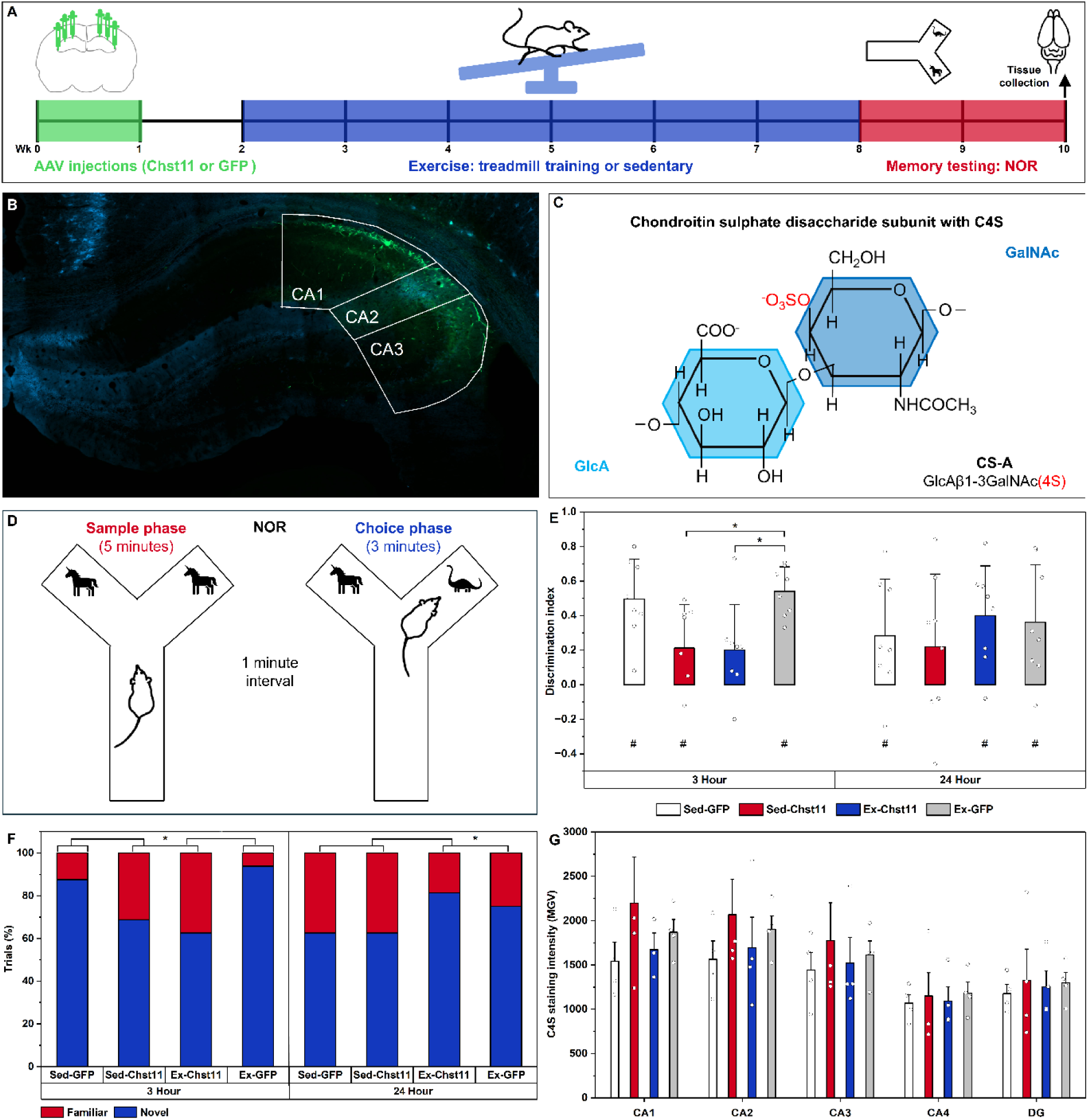
Exercise training attenuated memory impairments induced by *Chst11*. **A**. Study timeline. GFP: green fluorescent protein. NOR: Novel object recognition. **B**. GFP expression (green) in the CA1, CA2, and CA3 regions of the hippocampus which were targeted by AAV injections. Co-stained with *Wisteria floribunda* agglutinin (WFA) (blue). **C**. The chondroitin sulphate disaccharide units glucuronic acid (GlcA) and *N*-acetylgalactosamine (GalNAc) can be sulphated at several positions. CS-A is a disaccharide subunit which has GalNAc sulphation at position C4 (C4S). **D**. Schematic of the novel object recognition memory testing paradigm. **E**. Novel object recognition (NOR) results at three- and 24-hour time delays. Statistical significance determined using a one-way ANOVA for each timepoint. * *p*<0.05 when compared to Ex-GFP control. Hippocampal *Chst11* overexpression impaired NOR in both Sed-Chst11 (*p*=0.044) and Ex-Chst11 (*p*=0.006) animals at the 3-hour timepoint. One-sample t tests were used to determine whether groups could successfully perform NOR against chance (# - *p*<0.05). Bars represent mean discrimination index. Error bars: SE. *n*=8. **F.** Stacked bars present the mean percentage of NOR trials in which animals preferred either the novel or familiar objects following delays of either 3 hours or 24 hours. The percentage of trials with preference for the novel and familiar objects were compared between groups with a Chi-Squared test (χ2), * *p*<0.05, *n*=8. Both Chst11 groups showed a reduction in the percentage of trials in which animals correctly explored the novel object more than the familiar object at 3 hours (*p*<0.00001). At 24 hours, both exercise groups showed an increased percentage of trials successfully preferring the novel object (*p*=0.009). **G**. C4S staining intensity. MGV: mean grey value. Kruskal-Wallis H tests showed there was no statistically significant difference in any of the hippocampal subfields. DG – dentate gyrus. Bars represent mean MGV. Error bars: SE. *n*=4.

Therefore, instead of removing CS-GAGs, we aimed to use a viral vector to overexpress *Chst11* (a chondroitin-4-sulfotransferase) to upregulate C4S on CS-GAGs rendering them more inhibitory of neuroplasticity and thus impairing memory. We hypothesised that exercise training would downregulate the CS-GAGs which had increased levels of C4S, hence alleviating the memory impairments induced by overexpression of *Chst11*.

A study timeline is presented in (Figure 3A). We injected AAV1-PGK-CHST11 into the hippocampus of 16 rats to overexpress *Chst11* and therefore upregulate C4S, whilst 16 rats received control injections of AAV1-PGK-GFP (Figure 3B). From each group, eight rats remained sedentary (Sed-GFP and Sed-Chst11) whilst eight rats underwent a six-week programme of treadmill training (Ex-GFP and Ex-Chst11). At the end of the exercise programme, all rats were tested using the memory task, novel object recognition (Figure 3D), at both three- and 24-hour time delays. At the three-hour time delay, all experimental groups apart from Ex-Chst11 could successfully perform the object recognition task against chance (Sed-GFP: 0.495 ± 0.233 discrimination index, t_(7)_=6.011, *p*<0.001; Sed-Chst11: 0.213 ± 0.249 discrimination index, t_(7)_=2.415, *p*=0.046; Ex-GFP: 0.541 ± 0.141 discrimination index, t_(7)_=10.858, *p*<0.001; Ex-Chst11: 0.200 ± 0.264 discrimination index, t_(7)_=2.151, p=0.068) (Figure 3E). However, overexpressing *Chst11* impaired object recognition memory after the three-hour delay in both sedentary and exercise groups in comparison to Ex-GFP controls (Sed-Chst11: *p*=0.044; Ex-Chst11: *p*=0.034; F_(3,28)_=5.092, *p*=0.006). When using a 24-hour delay object recognition test, both Sed-GFP and Ex-GFP animals showed an overnight decline in memory, though both groups could successfully perform the task against chance (Sed-GFP: 0.281 ± 0.328 discrimination index, t_(7)_=2.424, *p*=0.046; Ex-GFP: 0.362 ± 0.338 discrimination index, t_(7)_=3.055, p=0.018). Although memory performance of Sed-Chst11 animals was comparable between the three- and 24-hour delays, they could no longer successfully perform the task against chance after the 24-hour delay (Sed-Chst11: 0.2205 ± 0.421 discrimination index, t_(7)_=1.481, *p*=0.182). However, whilst Ex-Chst11 animals could not successfully perform the memory task against chance after the three-hour delay, they were the only group to show improved overnight memory performance in the 24-hour delay memory task. This supports that exercise training mitigated the memory-impairment induced by overexpressing hippocampal *Chst11* as they could successfully perform the memory task against chance by the 24-hour time point (Ex-Chst11: 0.401 ± 0.286 discrimination index, t_(7)_=3.962, *p*=0.005).

In addition to analysing the discrimination index for object recognition memory performance, the percentage of trials in which animals correctly distinguished the novel objects was calculated (Figure 3F). Similar to the discrimination index, there was reduced novel object preference in Sed-Chst11 and Ex-Chst11 groups in comparison to GFP control groups at the 3-hour time delay (χ^2^= 39.17, *p*<0.00001). However, at the 24-hour time delay, both Ex-GFP and Ex-Chst11 groups showed improved performance of correctly distinguishing the novel objects compared to the Sed-GFP and Sed-Chst11 groups (χ^2^= 11.68, *p*=0.009). This further demonstrates that exercise training is capable of attenuating object recognition memory deficits induced by hippocampal *Chst11* overexpression at the 24-hour delay.

As the *Chst11* gene upregulates C4S, brain tissue was stained with an anti-C4S antibody. There was no statistically significant difference in hippocampal C4S staining intensity between any of the experimental groups (CA1: H_(3)_=1.842, *p*=0.606; CA2: H_(3)_=2.824, *p*=0.420; CA3: H_(3)_=0.728, *p*=0.867; CA4: H_(3)_=0.772, *p*=0.856; DG: H_(3)_=0.816, *p*=0.846) (Figure 3G).

However, the Sed-Chst11 group showed increased C4S staining compared to the Sed-GFP control group in the hippocampal regions where the viral injections were targeted (Sed-Chst11 C4S increase in CA1: 48.3 %, CA2: 37.8 %, and CA3: 29.5 % compared to Sed-GFP). The increase of C4S in the animals that received viral overexpression of *Chst11* was reduced in animals that underwent the six weeks of exercise training (Ex-Chst11 C4S reduction in CA1: -16.3 %, CA2: -8.7%, CA3: -10.5 % compared to Sed-Chst11). Together, these results support our hypothesis that the viral overexpression of *Chst11* would upregulate C4S, and that this would be alleviated by exercise training.

### PNNs of the hippocampal memory network

In the previous cohort which showed the reduction of CS-GAGs in PNNs within the CA1 and CA4 regions, brains were sampled directly at the end of the six-week exercise training programme. We hypothesised that the reduction of CS-GAGs in hippocampal PNNs would enhance neuroplasticity, thus making the hippocampal circuitry more flexible for integrating new memories. For the Chst11-memory cohort, animals underwent the six-week exercise training programme, after which they performed a series of novel object recognition tests throughout a two-week period in which all animals remained sedentary, and then brains were sampled following behavioural testing. PNNs have been documented to be modulated by behavioural testing (Carulli et al., 2020). Therefore, our experimental paradigm implemented exercise training first to prime the hippocampal circuitry and enhance neuroplasticity, and behavioural testing was performed after the cessation of exercise to investigate the hippocampal network after being exposed to new memory stimuli.

Consistent with results from the previous exercise cohort, the staining intensity of aggrecan and WFA was comparable between all experimental groups (Extended Data Figure 4). PNNs primarily enwrap parvalbumin positive interneurons and it has been documented that low parvalbumin network configurations are associated with enhanced structural synaptic plasticity and memory processes, whilst high parvalbumin network configurations induced the opposite (Donato et al., 2013). In support of exercise training enhancing hippocampal neuroplasticity, parvalbumin staining intensity was lower in all hippocampal subfields following exercise training, suggesting that the network was in a higher plasticity state (Figure 4A-C). However, the statistical analysis for parvalbumin staining intensity did not significantly differ between experimental groups (CA1: F_(3,12)_=1.171, *p*=0.361; CA2: F_(3,12)_=1.591, *p*=0.243; CA3: F_(3,11)_=0.895, *p*=0.474; CA4: F_(3,12)_=0.896, *p*=0.471; DG: F_(3,12)_=0.738, *p*=0.549).

**Figure 4.**
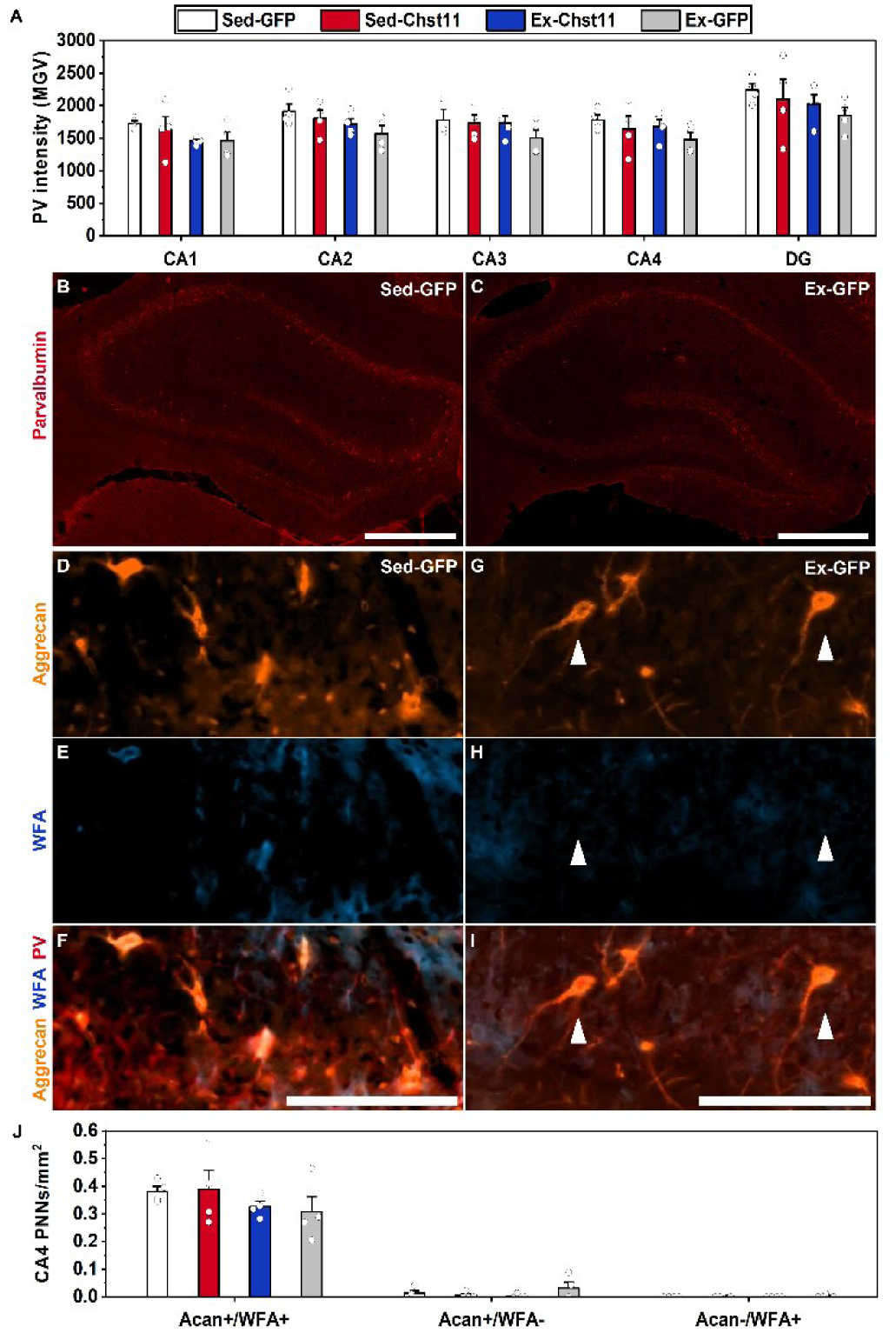
Hippocampal parvalbumin staining intensity and perineuronal nets. **A**. Parvalbumin (PV) staining intensity. MGV: Mean grey value. One-way ANOVAs showed there was no statistically significant difference in any of the hippocampal subfields. Though Parvalbumin staining was the lowest in Ex-GFP animals in all hippocampal subfields. DG – dentate gyrus. Bars represent mean MGV. Error bars: SE. *n*=4.**B-C**. Parvalbumin staining in the hippocampus of Sed-GFP and Ex-GFP animals, respectively. Scale bars: 200 μm. **D-I** Perineuronal nets (PNNs) labelled with aggrecan (**D** and **G**), *Wisteria floribunda* agglutinin (WFA) (**E** and **H**), and a MERGE of PNNs labelled with aggrecan and WFA co-stained with parvalbumin (**F** and **I**) in CA4 region of Sed-GFP and Ex-GFP. White arrowheads indicate PNNs that express aggrecan without WFA following exercise training. Scale bars: 50 μm. **J**. Perineuronal nets (PNNs) were counted as one of three different phenotypes: containing aggrecan and WFA (ACAN+ WFA+), containing aggrecan only (ACAN+ WFA-), or containing WFA only (ACAN-WFA+). ACAN: aggrecan antibody which labelled the core protein. WFA: *Wisteria floribunda* agglutinin which labelled CS-GAG chains. Lack of statistical significance determined using one-way ANOVAs. Bars represent mean PNN count. Error bars: SE. *n*=4.

Whilst PNNs mainly surround parvalbumin interneurons, they have also been observed around excitatory hippocampal granule cells (Carstens et al., 2016, Sorg et al., 2016). For the Chst11-memory cohort, we co-stained the PNN markers with parvalbumin to restrict the PNN count analysis around parvalbumin positive cells only. The counts of the three different PNN phenotypes (1: ACAN+/ WFA+, 2: ACAN+/WFA-, and 3: ACAN-/WFA+) were comparable between experimental groups in all hippocampal regions (Extended Data Figure 5). It is important to note that brain tissue in the Chst11-memory cohort was sampled after the two weeks of memory testing in which all animals remained sedentary. In a study using the bacterial enzyme, chondroitinase ABC, to completely degrade PNNs, WFA staining intensity had returned to 30% within two weeks (Lensjø et al., 2017). Therefore, the effect of exercise on the biochemical composition of PNNs may have been dampened in the two weeks from the cessation of exercise. However, PNN count results in the CA4 region are consistent with the previous exercise cohort. In comparison to the Sed-GFP group, the Ex-GFP group showed lower levels of ACAN+/WFA+ PNNs and a transition towards more PNNs with the ACAN+/WFA- (aggrecan only) phenotype (Figure 4D-J). Additionally, the number of ACAN+/WFA+ PNNs in CA4 was highest in the Sed-Chst11 group which was reduced in the Ex-Chst11 group.

In summary, these results suggest that exercise training renders the hippocampus a low-parvalbumin, high-plasticity region, and highlight CA4 as a key region in which exercise targets the modulation of PNNs within the hippocampal network.

During PNN analysis, we also observed prominent PNNs around parvalbumin negative neurons in the stratum oriens layer of CA3 (Extended Data Figure 6). In the cortex, PNNs have been observed around somatostatin interneurons, and as these interneuron subtypes are also found in stratum oriens of CA3, it is possible that the PNNs also surround hippocampal somatostatin interneurons (Honoré et al., 2021, Berretta et al., 2015).

## Discussion

Although exercise-induced enhancements of neurogenesis and memory are usually attributed to increased expression of the neurotrophic factor, BDNF (Fernandes et al., 2020, Neeper et al., 1995), the mechanism of exercise-induced neuroplasticity is still poorly understood. Central to our approach to tackle this was the notion that in addition to the parameters that enhance neuroplasticity, components of the system may also need to be downregulated to make way for neural changes. We implemented an exploratory approach to assess whether exercise training downregulated mRNA expression of genes within the RhoA/ROCK pathway, which is known to restrict neuroplasticity. We have shown that six weeks of moderate intensity treadmill training downregulated *Acan* in the hippocampus. *Acan* is a gene that encodes the CSPG, aggrecan, which is expressed in the loose extracellular matrix and is a main component of PNNs. We visualised PNNs in the hippocampus by labelling two different CSPG components: the aggrecan core protein and the CS-GAG chains which attach to the core protein. Exercise training induced a transition to a higher number of PNNs containing only aggrecan without labelling of CS-GAGs in the CA1 and CA4 regions of the hippocampus. As CS-GAGs are inhibitory to neuroplasticity and memory, we hypothesised that the exercise-induced reduction in CS-GAGs would enhance neuroplasticity and support improved memory. To test this, we designed a viral vector to overexpress *Chst11,* which adds C4S to chondroitin sulphate disaccharide units. C4S has been documented to restrict plasticity and impair memory (Wang et al., 2008, Yang et al., 2017). We hypothesised that the overexpression of *Chst11* in the hippocampus would upregulate chondroitin-4-sulphation and impair hippocampal dependent memory. We further hypothesised that exercise training would reduce levels of CS-GAGs which contain the upregulated chondroitin-4-sulphation in the hippocampus, and therefore attenuate memory impairments induced by *Chst11* overexpression. Our data show that the viral overexpression of *Chst11* in the hippocampus impaired novel object recognition memory. At the 24-hour time delay, sedentary animals which received hippocampal *Chst11* injections could not perform the object recognition tests successfully against chance. However, the animals that received hippocampal *Chst11* injections and underwent six weeks of exercise training could successfully perform the object recognition tests against chance, demonstrating that exercise training attenuated the memory deficits induced by *Chst11.* Collectively, our data support a mechanistic narrative that exercise training contributes to enhanced hippocampal neuroplasticity via the modulation of CS-GAGs in PNNs, which plays a role in memory performance.

In the hippocampus of mice housed with access to running wheels, PNNs were dynamically regulated throughout the diurnal 24-hour cycle; specifically, WFA+ PNNs (which were labelled for CS-GAGs) were downregulated during sleep and upregulated during wakefulness (Pantazopoulos et al., 2020). It is possible that the modification of CS-GAGs during sleep facilitates the consolidation of memories that were previously encoded that day. This supports our behavioural data in which exercise training improved object recognition memory performance after the 24-hour time delay and not after 3 hours. To further explore this mechanism, it would be advantageous to investigate how exercise-regulated diurnal rhythms of CS-GAG expression are associated with memory performance.

PNNs are expressed throughout the central nervous system and are implicated in numerous memory systems including cerebellar motor memory, fear memory, perirhinal cortex and object recognition memory, hippocampal spatial memory, and the hippocampal CA2 region and social memory (Rowlands et al., 2018, Carulli et al., 2020, Romberg et al., 2013, Gogolla et al., 2009, Huang et al., 2023, Maheu et al., 2025). However, this study demonstrated for the first time that both the modulation of CS-GAGs and chondroitin-4-sulphation in the hippocampus play a role in object recognition memory. Further supporting the downregulation of CS-GAGs for enhanced cognition, reduced levels of CS-GAGs observed in the frontal cortex of post-mortem human tissue were associated with resilience to cognitive impairment in subjects with Alzheimer’s disease pathology (de Vries et al., 2025). There is growing interest in chondroitin sulphate patterns: whilst *Chst11* and C4S are inhibitory to neuroplasticity, the sulphotransferase *Chst3* and C6S promote plasticity in the CNS (Miyata et al., 2012, Foscarin et al., 2017, Yang et al., 2021). Blocking C4S with an antibody alleviated memory deficits in a tauopathy mouse model (Yang et al., 2017). In contrast, C4S has been shown to prevent maladaptive PNN development in the CA2 region and is critical for social memory function (Huang et al., 2023). Furthermore, the overexpression of *Chst3* to upregulate the plasticity promoting C6S in the perirhinal cortex attenuated age-related object recognition memory impairment (Yang et al., 2021). This body of research highlights that manipulation of chondroitin sulphate patterns is a potent way of improving memory and provides a potential avenue of pharmaceutical development for regulating cognition.

We also observed PNNs surrounding parvalbumin-negative neurons in the stratum oriens layer of CA3 and hypothesise that these are PNN baring somatostatin interneurons. Future work should investigate the role of these PNNs within the hippocampal circuitry to elucidate the functional heterogeneity of PNNs in different regions and neuronal subtypes.

We have previously shown that voluntary wheel running reduces the number of PNNs in the rat hippocampus (Smith et al., 2015). Within voluntary wheel paradigms, Wistar rats can run for more than 250 minutes and cover up to 18 km per day (Ruegsegger et al., 2017). Such extensive levels of exercise are rare in humans; therefore, being able to modulate PNN composition with shorter bouts of moderate intensity treadmill training, as shown in this study, offers a more translatable and achievable approach to improving human health. Two recent studies in mice have also shown that exercise training modulates hippocampal PNNs labelled with WFA. Voluntary wheel running reduced the number of CA1 PNNs, which were negatively correlated with the number of neurogenic cells (Terstege et al., 2024). Additionally, treadmill training reduced PNNs in the mouse DG which was associated with improved spatial memory (Maheu et al., 2025). Whilst these studies support our results of exercise-induced modulation of PNNs, other work fails to elaborate how this impacts wider hippocampal circuitry for memory processing.

The hippocampus receives input from the medial and lateral entorhinal cortices which process spatial information; and the integration of non-spatial sensory information, respectively (Woods et al., 2018). In the trisynaptic pathway: 1) neurons from the entorhinal cortex project to granule cells in the DG via the perforant pathway; 2) DG mossy fibre projections contact CA3 pyramidal neurons; and 3) CA3 pyramidal neurons project to CA1 pyramidal cells via the Schaffer collateral pathway (Basu and Siegelbaum, 2015). Additionally, neurons from the entorhinal cortex project directly to the CA1 pyramidal neurons, bypassing the trisynaptic pathway. It is well established that exercise training increases neurogenesis (van Praag et al., 1999). Neurogenic adult-born granule cells originate in the subgranular zone of the DG and their dendritic trees develop out into the molecular layer where they receive input from the entorhinal cortex via the perforant path (Toni and Schinder, 2016). Adult-born granule cell mossy fibre boutons are documented to contact parvalbumin interneurons enwrapped by PNNs between the two blades of the DG, (Briones et al., 2021) and to pyramidal and parvalbumin neurons in the CA3 region (Toni and Schinder, 2016). Our results show that exercise training modulates PNNs in both CA1 and CA4 in the rat hippocampus (what we refer to as the CA4 region is the pyramidal cell layer that terminates between the blades of the DG). As PNNs are structures that restrict synaptic remodelling, we propose that the exercise-induced modulation of CS-GAGs in the CA4 region makes PNNs more plastic and supports the integration of neurogenic adult-born granule cells into the existing hippocampal circuitry, both in the CA4 region and downstream the trisynaptic pathway at CA1. In support of this, when we manipulated CS-GAGs to be more inhibitory via overexpressing *Chst11* in the hippocampus, novel object recognition memory was impaired. It is plausible that the overexpression of *Chst11* in the CA1 and CA3 regions made it more difficult for adult-born granule cells to integrate within the existing hippocampal circuitry and therefore interrupted the flow of new memory processing throughout the trisynaptic circuit. In support of this, the application of synthetic chondroitin sulphate with a high C4S:C6S ratio reduced synaptic transmission in the DG and CA1 of hippocampal slices (Albiñana et al., 2015).

In summary, we have shown that exercise training modulates *Acan* expression, alters the biochemical composition of PNNs, and mitigates memory deficits induced by overexpression of hippocampal *Chst11.* Our results support that in the fields of exercise-induced neuroplasticity and for enhancing memory and learning, the modulation of inhibitory pathways/molecules that restrict plasticity is an important piece of the puzzle that should be considered by future researchers. To our knowledge, we are the first to propose a mechanistic narrative in which exercise training and the modulation of CS-GAGs/PNNs enhance neuroplasticity within the hippocampal network to potentially support the integration of neurogenic adult-born granule cells for memory function. This is a vital step towards understanding how exercise improves memory which is fundamental for human health and for conditions in which cognitive decline is prominent. These results propel our current understanding of how exercise can be harnessed to augment neuroplasticity and provides a new translational direction for exercise mimetic and drug discovery research aiming to replicate the beneficial effects of exercise.

## Methods

### Animals

Adult male Wistar rats (200-250 g) were obtained from Central Biomedical Services (University of Leeds, UK, *n*=6) and Charles River Laboratories (Canterbury, UK, *n*=40). Animals were housed four to a cage with *ad libitum* access to food and water, at 20 ± 1 °C, and under a 12-hour light/dark cycle at Central Biomedical Services (University of Leeds, UK). All procedures were conducted during the light cycle, complied with the UK Animals (Scientific Procedures) Act 1986 (ASPA), and were approved by the University of Leeds Animal Welfare and Ethical Review Committee.

### Exercise training protocol

Rats were habituated to a motorised treadmill (Panlab, Harvard apparatus, UK) for five minutes over five consecutive days. Throughout the five days, the treadmill speed was gradually increased from 0 cm.s^-1^ to 50 cm.s^-1^ (Supplementary Table 1). If animals turned to face the back of the treadmill, the speed was lowered until the animal returned to forward facing (usually within 30 seconds) and then returned to the required speed.

For the exercise training protocol, animals ran for a total of 33 minutes on the treadmill at 5° inclination, five days a week, for six weeks. The first three minutes of the protocol served as a warmup at 25 cm.s^-1^. Following the warmup, animals ran for 30 minutes at 32 cm.s^-1^ (approximately 67% VO_2 max_; (Qin et al., 2020). The control group were brought into the behavioural room during training times though remained sedentary throughout the six-week training period.

### Experimental cohorts

For cohort 1 (Exercise cohort), a total of 14 rats were used (Sedentary *n*=7, Exercise *n*=7). Animals underwent their experimental conditions (sedentary or exercise) for six weeks and brain samples were taken 24 hours after the final experimental day. Four animals per group were used for gene expression experiments and three animals per group were used for immunohistochemistry. For cohort 2 (Chst11 memory cohort) a total of 32 rats were used (Sedentary-GFP *n*=8, Sedentary-Chst11 *n*=8, Exercise-GFP *n*=8, Exercise-Chst11 *n*=8). Animals received hippocampal stereotaxic injections of AAVs to express either *Chst11* or GFP. Animals underwent their experimental conditions (sedentary or exercise) for six weeks. After the training protocol, all 32 animals were used for behavioural memory testing. Brain samples were taken 24 hours after the final day of memory testing and four animals per group were used for immunohistochemistry.

### Reverse transcription quantitative polymerase chain reaction (RT-qPCR)

Animals were overdosed with an intraperitoneal injection of sodium pentobarbital (200 mg/kg) to induce deep anaesthesia 24 hours after the exercise training protocol was over. Once the blink and pedal reflexes were absent, rats were decapitated and the hippocampus, cortex (directly above the hippocampus), and lumbar region of the spinal cord were snap frozen using liquid nitrogen and stored at −80°C. Gene expression experiments were informed by the MIQE guidelines (Bustin et al., 2009) and were performed as described in (Doody et al., 2024). Tissue samples were homogenised in 700 µL of TRIzol® using 7 mm stainless steel ball bearings and a TissueLyser LT bead mill (Qiagen, Germany) for 10 minutes. RNA was extracted using TRIzol® and the PureLink® RNA Mini Kit with On-column PureLink® DNase Treatment (ThermoFisher Scientific, UK). RNA concentration and purity were determined with a NanoDrop™ ND-2000 spectrophotometer. RNA was diluted to 200 ng/µl using ultra-pure nuclease free water. cDNA was reverse transcribed from 2 µg of RNA using the Precision nanoScript TM2 Reverse Transcription kit (PrimerDesign, UK). cDNA was diluted to 5 ng/µl using ultra-pure nuclease free water. Custom TaqMan® Array Plates were used to quantify the relative gene expression of 25 genes within the RhoA/ROCK pathway (Supplementary Table 2). Plates were briefly centrifuged before adding the cDNA-master mix. Each 20 µL reaction contained 10 µL of cDNA at 2 ng/µL (total cDNA per reaction: 20 ng) and 10 µL of TaqMan™ Fast Advanced Master Mix (ThermoFisher Scientific, UK). Plates were sealed with optical adhesive film, briefly centrifuged, and loaded on a CFX96 Touch™ Real-Time PCR Detection System (Bio-Rad, UK). The thermal protocol included a UNG incubation at 50°C for two minutes, an enzyme activation period at 95°C for 20 seconds, and 40 cycles of a one second denaturing step at 95°C and a 20 second anneal/extension step at 60°C. Samples were run in duplicate.

### RT-qPCR data processing

The relative gene expression of target genes was calculated using the 2^-ΔΔCT^ method. Relative gene expression was normalised using the reference genes 18s and Cyclophilin A. Data were excluded if outside two standard deviations from the mean or if the Cq value was above cycle 35. Sample sizes for target genes after exclusions were applied are presented in Supplementary Table 3. Relative fold change cut off values of ≤0.75 and ≥1.5 were used to further explore the gene expression profiles of the RhoA/ROCK pathway (Kubo et al., 2015). The STRING database was used to generate a protein interaction network of the gene expression profiles created with the cut off values (Szklarczyk et al., 2023). The STRING enrichment analysis was aligned the gene list to KEGG pathways and Gene Ontology terms.

### Immunohistochemistry

Animals were overdosed with an intraperitoneal injection of sodium pentobarbital (200 mg/kg) to induce deep anaesthesia. Once the blink and pedal reflexes were absent, rats were transcardially perfused with sodium phosphate buffer (1 M PB), followed by 4% paraformaldehyde (PFA), and a final rinse with 1 M PB. Brains were post-fixed in 4% PFA overnight at 4°C. Tissue was cryoprotected in a 30% sucrose solution with 1 M PB and stored at 4°C. Brains were mounted in OCT (FSC 22 Frozen Section Media; Leica Biosystems), frozen with dry ice, and stored at −80°C. Samples were transferred to −20°C overnight to equilibrate to the temperature used within the cryostat chamber. Tissue was cut into 25 µm coronal sections using a cryostat (Leica CM1850; Leica Biosystems). Sections were collected into 1 M PBS before being transferred into cryoprotectant and stored at −20°C.

Free-floating sections were washed at room temperature, for three five-minute periods in 1 M PBS to remove the cryoprotectant. The sections were blocked for two hours in 0.2% PBST (1 M PBS and 0.2% Triton X-100) and 3% normal donkey serum (NDS). The sections were then incubated overnight in 0.2% PBST and 3% NDS, containing the primary antibodies at 4°C. Primary antibodies used were anti-NeuN (MAB377, 1:500, Mouse, Merck Millipore, Lot:3832727, RRID:AB_2298772), anti-Aggrecan (#AB1031, 1:250, Rabbit, Merck Millipore, Lot:4206909, RRID:AB_90460), biotin-conjugated *Wisteria floribunda* agglutinin (WFA) (L1516, 1:300, Sigma Aldrich, Lot:SLBS9583, RRID:AB_2620171), anti-Parvalbumin (PVG213, 1:10000, Goat, Swant, Lot:PVG13_210401_200, RRID:AB_2721207), anti-Chondroitin sulphate A (Clone 2H6) (NU-07-001, 1:50, Mouse IgM, CosmoBio, Lot: MA-007-01), anti-NeuN (ABN78, 1:500, Rabbit, Merck Millipore, Lot:3304299, RRID:AB_10807945). Sections were then washed three times for 10 minutes in 1 M PBS before being incubated for two hours at room temperature in 1 M PBS containing secondary antibodies. Secondary antibodies were: donkey anti-mouse (A21202, 1:1000, Alexa Fluor® 488, Invitrogen, Lot:2090565, RRID:AB_141607), donkey anti-rabbit (A31572, 1:1000, Alexa Fluor® 555, Invitrogen, Lot:2088692, RRID:AB_162543), streptavidin (S11222, 1:250, Pacific Blue™, Invitrogen, Lot:2260873), donkey anti-goat (A21447, 1:1000, Alexa Fluor® 657, Invitrogen, Lot:2674361, RRID:AB_2535864), donkey anti-mouse IgM µ chain specific (715-605-020, 1:250, Alexa Fluor® 647, Jackson Immuno, Lot: 153 845, RRID:AB_2340860). The tissue was then washed three times for 10 minutes in 1 M PBS and washed twice for five minutes in tris non-saline (1 M TNS). Sections were mounted onto microscope slides (Superfrost Plus™, ThermoFisher Scientific) using mounting medium (VECTASHIELD® Antifade Mounting Medium, Vector Laboratories) or FluorSave^TM^ Reagent (Merck, Lot:4226691). Negative control tissue was incubated with only secondary antibodies (no primary antibodies).

### Image acquisition

Images were acquired in the Bio-Imaging Facility at the University of Leeds. Full tile scans of the dorsal hippocampus (between Bregma -3.14 and -3.80 mm) were taken at 20X magnification on an Axio Scan Z1 slide scanner (Zeiss, UK). Images were assigned a new code so that image analysis could be performed blinded to experimental groupings.

### Image analysis

All image analysis was conducted in the software Fiji for ImageJ (Schindelin *et al*., 2012).. Tissue sections between Bregma -3.14 and -3.80 mm that displayed the dorsal hippocampus were chosen for analysis. Five sections were imaged per animal to ensure that a minimum of three sections could be analysed. All analysis was performed bilaterally. The results were normalised (as outlined below) and were then averaged between the two hemispheres. The mean of normalised results was calculated per animal (mean of 3-5 sections) and then per experimental group (mean of 3-4 animals). Regions of interest (ROIs) were manually drawn with the polygon tool covering the CA regions (1-4) and dentate gyrus of the hippocampus with reference to Bregma -3.30 mm in the Rat Brain Atlas (Paxinos and Watson, 2013) (Supplementary Figure 1).

For aggrecan, WFA, parvalbumin, and C4S, staining intensity (mean grey value) and area were recorded for each ROI. Staining intensity was normalised by subtracting the mean grey value for the respective negative control from the experimental mean grey value. For each ROI, the normalised staining intensity was averaged between the two hemispheres. The mean normalised staining intensity was then taken for all sections per animal and then for each animal per experimental group.

PNNs were counted using the Fiji Image J Cell Counter plugin. Three different PNN phenotypes were counted: 1) PNNs containing both aggrecan and WFA (ACAN+/WFA+); 2) PNNs containing only aggrecan (ACAN+/WFA-); and 3) PNNs containing only WFA (ACAN-/WFA+). PNN counts were normalised to the area of the ROI and converted from μm to mm. The total PNN count was calculated as the sum of all three phenotype counts. For the exercise cohort, all visible PNNs were counted. For the Chst11 memory cohort, PNNs were co-stained with parvalbumin, and only PNNs surrounding parvalbumin positive cells were counted.

PNN thickness was measured by drawing four intersecting lines for the outer distance of the PNN and four intersecting lines for the inner distance of the PNN. PNN thickness was calculated by subtracting the mean inner PNN distance from the mean outer PNN distance and dividing this value by 2. PNN thickness was analysed separately for aggrecan and WFA positive PNNs for each hippocampal region. All visible PNNs were included in analysis if the outer and inner edges of the PNN could be clearly defined.

### Stereotaxic surgeries

Animals were anaesthetised using isoflurane (5 % in O_2_ for induction and 2.5 % in O_2_ for maintenance). The dorsal surface of the head was shaved and sterilised (50% iodine, 50% ethanol solution), and eye lubricant (Lubrithal) was applied. Animals were placed on a homeothermic heat pad (Harvard Apparatus Homoeothermic Monitoring System; Harvard Apparatus, Massachusetts, USA) and were secured in a stereotaxic frame throughout the surgery. An incision was made down the midline of the head in the sagittal plane and connective tissues were pushed away with a cotton bud to expose landmarks of the skull. Bone wax was applied to the skull to suppress bone bleeding. The head was aligned so that DV readouts for Bregma and Lambda were within 0.1 mm of each other. Bilateral craniotomy windows (approximately 3.5 x 3 mm) were drilled using a dental drill and 1 mm round-headed drill bits (Saeshin Dental STRONG 90 Micro motor with 102S Hand piece 35,000 RPM; Saeshin Precision Co., Daegu, Korea). Skin retractors were used to keep the craniotomy windows clear. Six x 0.5 μL injections (3 each hemisphere into CA1, CA2, and CA3 - total virus volume 3 μL per rat) of either AAV1-PGK-GFP or AAV1-PGK-CHST11 (Vigene Biosciences, Maryland, USA) were injected into the dorsal hippocampus. All coordinates given in cm from Bregma (zero) (Left CA3: AP -0.33, ML -0.35, DV -0.36; Left CA2: AP -0.33, ML - 0.3, DV -0.3; Left CA1: AP -0.33, ML -0.2, DV -0.26; Right CA1: AP -0.33, ML 0.2, DV -0.26; Right CA2: AP -0.33, ML 0.3, DV -0.3; Right CA3: -0.33, ML 0.35, DV -0.36). The 0.5 μL of virus was injected over two minutes (250 nL/min) using a 5 μL Hamilton Syringe (26-gauge bevelled needle) and a UMP3 Microinjection Syringe Pump with Micro4 controller (WPI Instruments, Florida, USA). The needle was left in position for two minutes to allow residual AAV to be absorbed before being slowly withdrawn. After the final injection the skin was sutured with non-absorbable suture. Pain relief (Vetergesic Buprenorphine; 0.015 mg/kg; Henry Schein Animal Health, Dumfries, UK), antibiotics (Baytril enrofloxacin; 2.5 mg/kg; Henry Schein Animal Health, Dumfries, UK), were administered by subcutaneous injection before removal from anaesthesia and for three days post-surgery. Animals were allowed to recover in a heated chamber until fully conscious and then returned to their home cage.

### Behavioural testing – Novel object recognition

A Y-shaped apparatus was constructed with 10 mm matt white foam PVC (Cut to size, Sheet Plastics, UK) using dimensions from Winters et al. (2004). Novel object recognition experiments were performed as outlined in (Romberg et al., 2013). The arm at the base of the Y-shaped apparatus was closed off by a guillotine door where the rats were placed at the start of the experiments. The other two shorter arms contained objects which the rat could explore once the guillotine door was opened. All rats were familiarised to the Y-shaped apparatus that contained treats (mini marshmallows) in the shorter arms for five minutes over two consecutive days. In the sample phase of the experiments two identical objects were placed in the short arms which rats could explore for five minutes. Following a delay of either three or 24 hours, the rats returned to the apparatus for the choice phase. In the choice phase the rats explored a new pair of objects for three minutes: one identical to the sample object pair, and one completely novel object. The objects were held in place with white tack (UHU, Germany). Between each trial the objects and the apparatus were cleaned with fragrance free detergent (Citop Zero, Green Speed, Netherlands) to remove olfactory cues. The rats received two trials at each different delay timepoint (four trials in total) with 48 hours rest in between each trial. A different set of object pairings were used for each of the four trials (Supplementary Figure 2). Four different random orders of objects were used throughout the experiments. The choice phase was recorded using a camcorder and tripod above the Y-shaped apparatus. The time rats spent exploring the familiar and novel objects during the choice phase was recorded. Object exploration was timed when the rat was directly looking at/sniffing the object. Exploration was not timed if the rat looked away, stood above/on top of the object, or was chewing tack.

The discrimination index was calculated by subtracting the time spent exploring the familiar object from the time spent exploring the novel object, then dividing this value by the total exploration time. The discrimination index was calculated for one minute of video footage from the choice phase as the strongest novel object preference scores usually occur within the first minute of the test phase (Jablonski et al., 2013). The discrimination index gives a score between -1 and 1. A positive score shows that more time was spent exploring the novel object. Whereas a negative score shows that more time was spent exploring the familiar object. Rodents are expected to remember that they have already seen the familiar object and spend more time exploring the novel object. Thus, rats with memory impairments would be expected to have lower discrimination indices.

### Statistical analysis

Statistical analysis was performed using IBM SPSS Statistics Version 30.0 (RRID:SCR_002865). Normality was assessed with the Shapiro-Wilk test prior to conducting statistical tests. The alpha level was set at 0.05 and statistical significance was denoted in figures with an asterisk (*p<0.05). In the exercise cohort, independent t tests were used for comparing gene expression and immunohistochemical data between sedentary and exercise animals, unless data was not normally distributed in which Mann-Whitney U tests were used. In the Chst11 memory cohort, one sample t tests were used to determine whether each experimental group could successfully perform NOR tests against chance. One-way ANOVAs followed by Bonferroni post hoc tests were used to analyse the discrimination index from NOR tests. Chi-square (χ^2^) tests were used to compare the percentage of NOR trials in which animals preferred the novel and familiar objects. C4S staining intensity was analysed using Kruskal-Wallis H tests. One-way ANOVAs followed by Bonferroni post hoc tests were used to analyse staining intensity of aggrecan, WFA, and parvalbumin. Figures were created using OriginPro Version 2025b (RRID:SCR_014212).

## Author Contributions

**N.D, R.I, J.K,** and **G.A:** design and conceptualisation of the research. **N.D, B.S, N.S, V.L** and **E.A:** conducting experiments. **S.B** and **R.H:** fluorescent imaging. **N.D:** data collection, data analysis, statistical analysis, and manuscript/figure preparation. **N.D, R.I, G.A, B.S, N.S, V.L** and **E.A:** manuscript reviewing and editing.

---

**Extended Data Figure 5.**
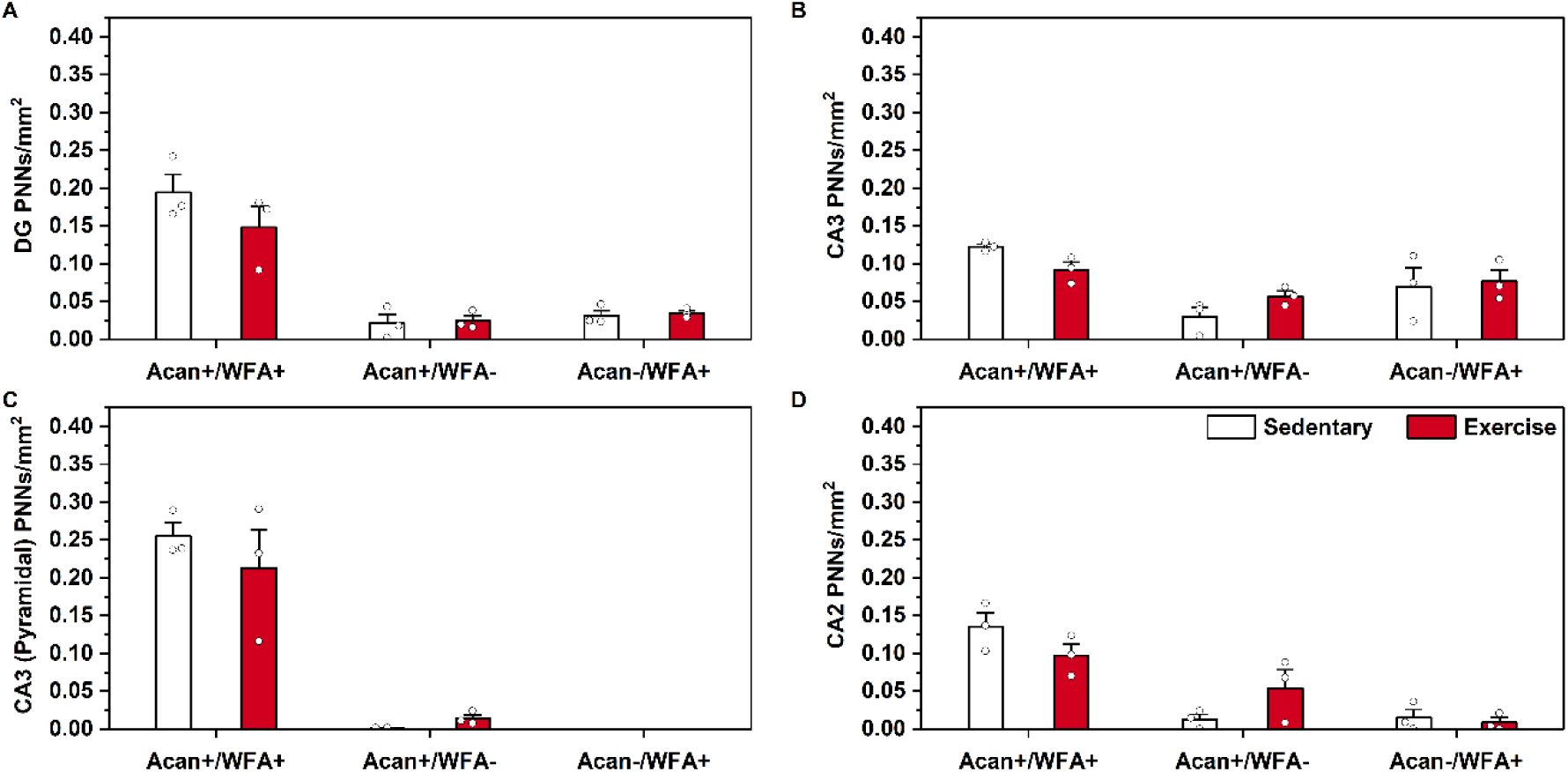
PNN phenotype counts in the hippocampus following exercise training. Perineuronal nets (PNNs) were counted as one of three different phenotypes: containing aggrecan and WFA (ACAN+ WFA+), containing aggrecan only (ACAN+ WFA-), or containing WFA only (ACAN-WFA+). ACAN: aggrecan antibody which labelled the core protein. WFA: *Wisteria floribunda* agglutinin which labelled chondroitin sulphate glycosaminoglycan (CS-GAG) chains. PNN counts for dentate gyrus (DG) (**A**), CA3 (**B**), CA3 pyramidal layer (**C**), and CA2 (**D**). Statistical significance was assessed using an independent t test between sedentary and exercise animals. None of the comparisons reached statistical significance. Bars represent mean PNN count. Error bars: SE. *n*=3.

**Extended Data Figure 6.**
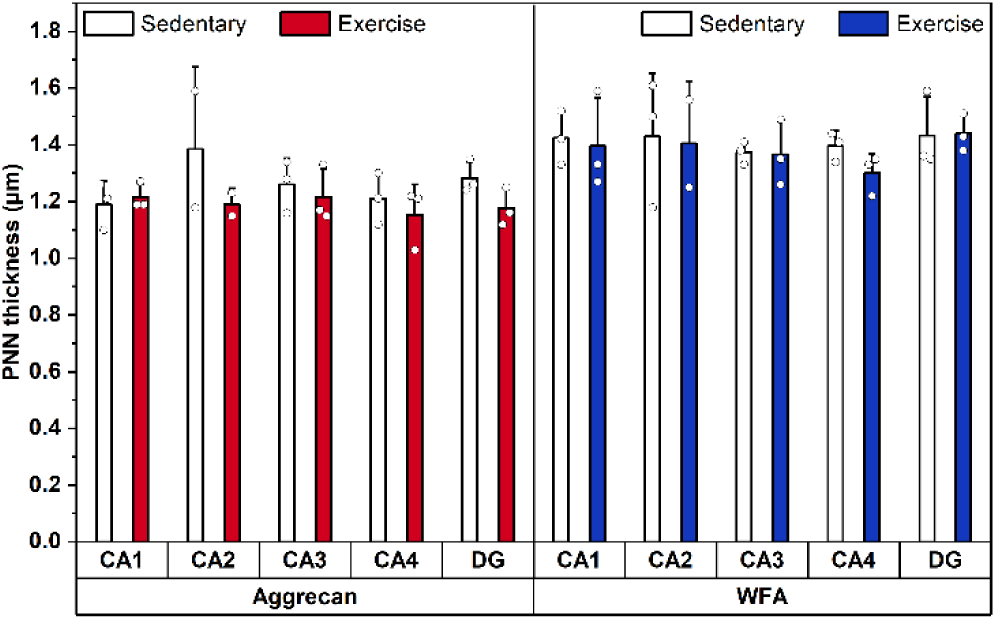
Hippocampal PNN thickness following exercise training. Perineuronal net (PNN) thickness was comparable between Sedentary and Exercise groups for both aggrecan and *Wisteria floribunda* agglutinin (WFA) labelled PNNs. Statistical significance was assessed using an independent t test between sedentary and exercise animals. None of the comparisons reached statistical significance. Bars represent mean PNN thickness. Error bars: SE. *n*=3.

**Extended Data Figure 7.**
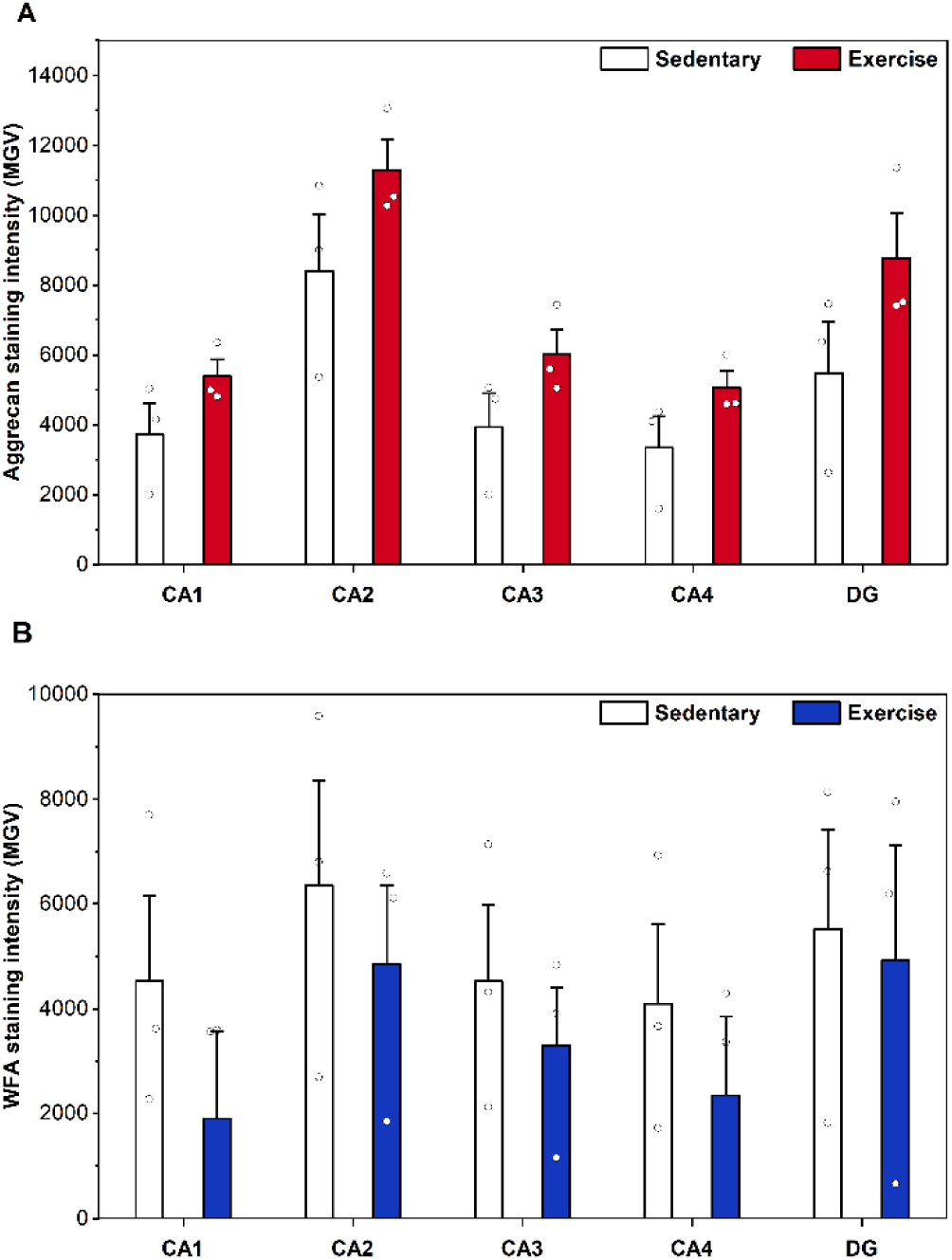
Hippocampal PNN staining intensity following exercise training. **A**. Aggrecan staining intensity. **B**. *Wisteria floribunda* agglutinin (WFA) staining intensity. MGV: mean grey value. Independent t tests showed there was no statistically significant difference between Sedentary and Exercise animals in any of the hippocampal subfields. DG – dentate gyrus. Bars represent mean MGV. Error bars: SE. *n*=3.

**Extended Data Figure 8.**
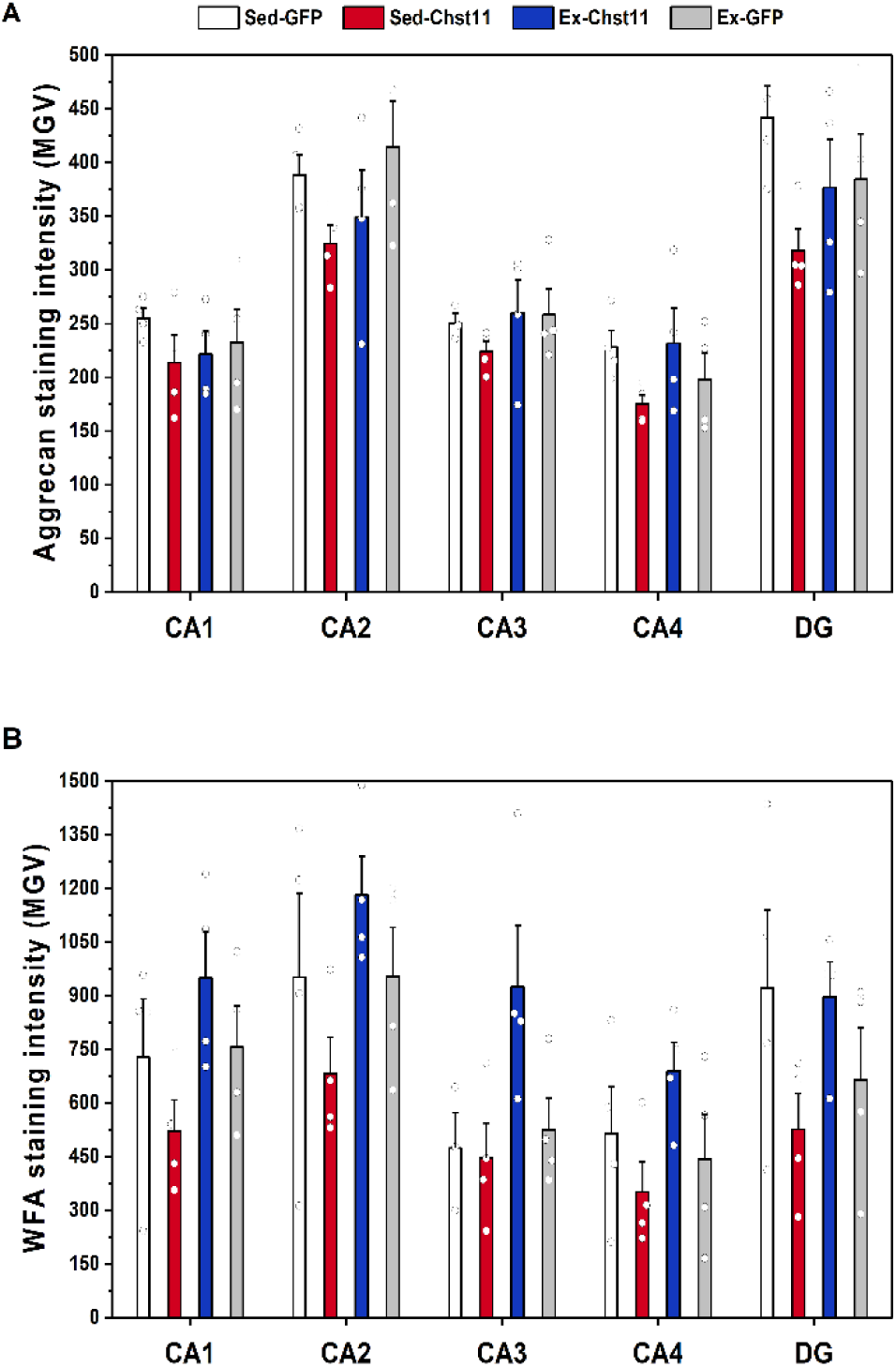
Hippocampal PNN staining intensity following *Chst11* overexpression and exercise training. **A**. Aggrecan staining intensity. **B**. *Wisteria floribunda* agglutinin (WFA) staining intensity. MGV: mean grey value. One-way ANOVAs showed there was no statistically significant difference between experimental groups in any of the hippocampal subfields. DG – dentate gyrus. Bars represent mean MGV. Error bars: SE. *n*=4.

**Extended Data Figure 9.**
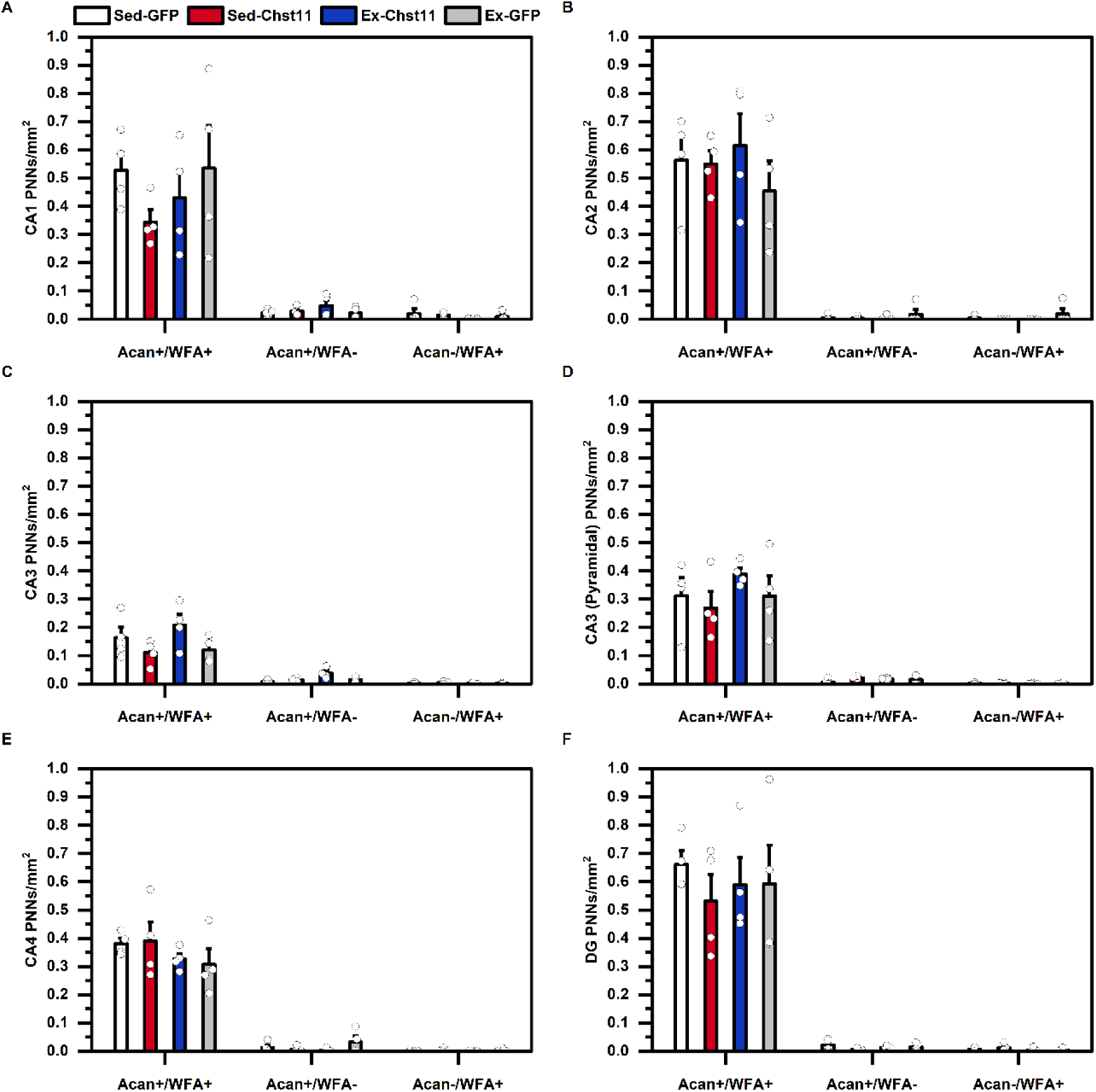
Hippocampal PNN phenotype counts following *Chst11* overexpression and exercise training. Perineuronal nets (PNNs) were counted as one of three different phenotypes: labelled with aggrecan and WFA (ACAN+ WFA+), labelled with aggrecan only (ACAN+ WFA-), or labelled with WFA only (ACAN-WFA+). ACAN: aggrecan antibody which labelled the core protein. WFA: *Wisteria floribunda* agglutinin which labelled chondroitin sulphate glycosaminoglycan (CS-GAG) chains. PNN counts for CA1 (**A**), CA2 (**B**), CA3 (**C**), CA3 pyramidal layer (**D**), CA4 (**E**), and dentate gyrus (DG) (**F**). Statistical significance determined using one-way ANOVAs. All results not statistically significant. Bars represent mean PNN count. Error bars: SE. *n*=4.

**Extended Data Figure 10.**
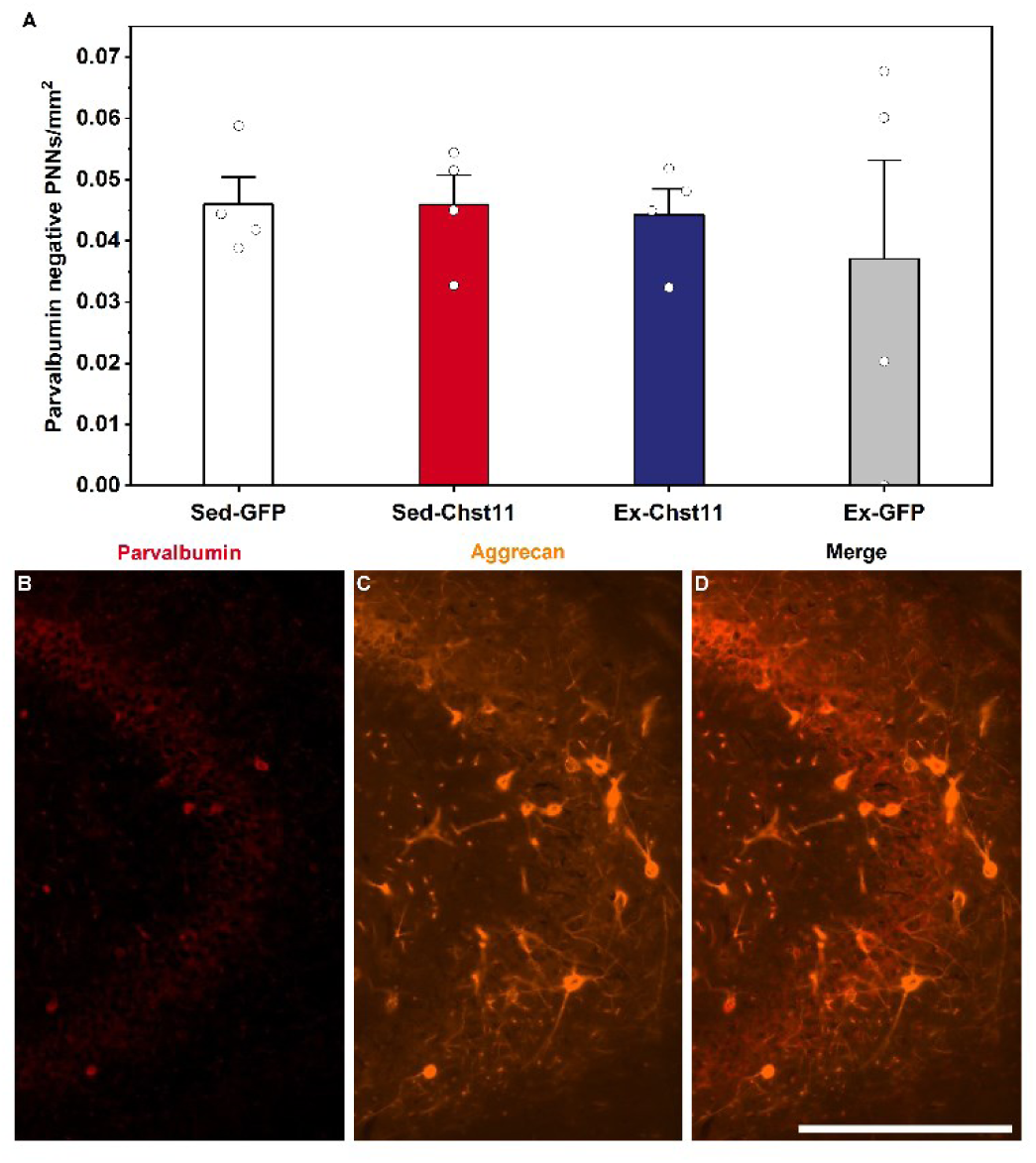
Parvalbumin negative PNNs observed in the hippocampus. **A**. Number of parvalbumin negative perineuronal nets (PNNs) observed in the stratum oriens layer of CA3. Statistical significance determined using one-way ANOVAs. All results not statistically significant. Bars represent mean PNN count. Error bars: SE. n=4. **B-D** Images showing aggrecan positive PNNs that are not surrounding parvalbumin cells in the stratum oriens layer of CA3. Scale bar: 100 µm.

**Extended Data Table 1.** Gene expression of RhoA/ROCK pathway components following exercise training.

| Mean relative gene expression (Log2 FC) compared to sedentary control (±SD) |  |  |  |  |  |  |
| --- | --- | --- | --- | --- | --- | --- |
| Target gene | Hippocampus | Statistical analysis | Cortex | Statistical analysis | Lumbar | Statistical analysis |
| ACAN | -0.6078 (0.1719) | $t_{(4)} = 2.880, p = 0.045$ | -0.1221 (0.1988) | - | 0.1988 (0.6642) | $t_{(6)} = -0.629, p = 0.552$ |
| BCAN | -0.4544 (0.7611) | $t_{(6)} = 0.822, p = 0.443$ | 0.4396 (0.0893) | $t_{(5)} = -0.796, p = 0.462$ | -0.3315 (0.7097) | $U = 4.000, p = 0.343$ |
| CFL1 | -0.0126 (0.4561) | $t_{(6)} = 0.029, p = 0.977$ | -0.3634 (0.5160) | $t_{(5)} = 0.936, p = 0.392$ | 0.0707 (0.5885) | $U = 8.000, p = 1.000$ |
| DPYSL2 | -0.5966 (0.4854) | $U = 3.000, p = 0.200$ | 0.4002 (0.4463) | $t_{(5)} = -1.162, p = 0.298$ | 0.0227 (0.4770) | $t_{(6)} = -0.081, p = 0.938$ |
| EFNB3 | 0.1808 (0.4605) | $t_{(6)} = -0.458, p = 0.663$ | - | - | - | - |
| GNA13 | -0.6927 (0.7413) | $U = 2.000, p = 0.114$ | 0.1923 (0.1035) | $t_{(5)} = -1.088, p = 0.326$ | 0.1851 (0.3887) | $t_{(6)} = -0.736, p = 0.489$ |
| LARG | -0.2157 (0.4591) | $t_{(6)} = 0.807, p = 0.451$ | -0.1285 (0.1259) | $t_{(5)} = 0.705, p = 0.512$ | -0.1757 (0.2410) | $t_{(6)} = 0.612, p = 0.563$ |
| LIMK1 | -0.9166 (0.0388) | - | 0.0530 (0.3094) | $U = 7.000, p = 1.000$ | -0.1771 (0.2710) | $t_{(6)} = 0.752, p = 0.480$ |
| LINGO-1 | -0.4201 (0.6351) | $t_{(6)} = 1.242, p = 0.261$ | -0.1354 (0.2545) | $t_{(5)} = 0.942, p = 0.390$ | -0.0050 (0.2534) | $t_{(6)} = 0.035, p = 0.973$ |
| LPAR1 | 0.1274 (0.7306) | $t_{(6)} = -0.301, p = 0.774$ | -0.0523 (0.1366) | $t_{(5)} = 0.216, p = 0.838$ | 0.1040 (0.2198) | $t_{(6)} = -0.641, p = 0.545$ |
| MAG | 0.2420 (0.7568) | $t_{(6)} = -0.607, p = 0.566$ | 0.2540 (0.2305) | $t_{(5)} = -0.704, p = 0.513$ | 0.1984 (0.5661) | $t_{(6)} = -0.650, p = 0.540$ |
| MYL2 | - | - | - | - | 0.0334 (0.5203) | $t_{(5)} = -0.096, p = 0.927$ |
| NCAN | -0.3074 (0.4003) | $U = 6.000, p = 0.686$ | 0.1465 (0.1887) | $U = 9.000, p = 0.400$ | 0.1667 (0.4080) | $t_{(6)} = -0.653, p = 0.538$ |
| NgR1 (Rtn4r) | -0.3368 (0.7758) | $t_{(6)} = 0.584, p = 0.580$ | 0.3705 (0.2665) | $t_{(5)} = -0.978, p = 0.373$ | -0.3152 (0.8212) | $t_{(6)} = 0.741, p = 0.487$ |
| NgR2 (Rtn4rl2) | -0.8084 (0.0950) | - | 0.0531 (0.3369) | $t_{(5)} = -0.144, p = 0.892$ | - | - |
| NgR3 (Rtn4rl1) | -0.8081 (0.2445) | $t_{(6)} = 2.154, p = 0.084$ | 0.0388 (0.1705) | $t_{(5)} = -0.141, p = 0.895$ | - | - |
| Nogo-A (Rtn4) | 0.3194 (0.3726) | $t_{(6)} = -1.260, p = 0.245$ | 0.0103 (0.3618) | $t_{(5)} = -0.043, p = 0.968$ | -0.0275 (0.2194) | $t_{(6)} = 0.137, p = 0.896$ |
| OMG | -0.9671 (1.0012) | $t_{(6)} = 1.858, p = 0.112$ | -0.3484 (0.1176) | $t_{(5)} = 3.622, p = 0.015$ | -0.0977 (0.2904) | $t_{(6)} = 0.547, p = 0.604$ |
| PTEN | 0.0249 (0.5664) | $t_{(6)} = -0.078, p = 0.940$ | 0.0524 (0.6509) | $t_{(5)} = -0.125, p = 0.906$ | -0.0271 (0.3572) | $t_{(6)} = 0.071, p = 0.946$ |
| PTPRF | 0.4256 (0.9082) | - | -0.1568 (0.1609) | - | -0.2278 (0.0391) | $t_{(4)} = 2.311, p = 0.082$ |
| PTPRS | - | - | -0.4806 (0.0250) | $t_{(4)} = 2.687, p = 0.055$ | - | - |
| ROCK2 | -0.9755 (0.9308) | $t_{(6)} = 2.042, p = 0.087$ | -0.1745 (0.1367) | $t_{(5)} = 1.101, p = 0.321$ | -0.0879 (0.1210) | $t_{(6)} = 0.543, p = 0.607$ |
| SEMA4D | 0.2554 (0.5620) | $t_{(6)} = -0.569, p = 0.590$ | 0.4053 (0.3918) | $t_{(5)} = -1.053, p = 0.340$ | -0.4414 (0.8720) | $t_{(6)} = 0.640, p = 0.546$ |
| TROY (Tnfrsf19) | -0.1514 (0.6697) | $t_{(6)} = 0.420, p = 0.689$ | -0.1767 (0.1561) | $t_{(5)} = 1.442, p = 0.209$ | 0.0227 (0.2967) | $t_{(6)} = -0.077, p = 0.941$ |
| VCAN | 0.1001 (0.4484) | $t_{(6)} = -0.391, p = 0.709$ | -0.1863 (0.1370) | $U = 2.000, p = 0.229$ | -0.0718 (0.2650) | $t_{(6)} = 0.328, p = 0.754$ |

**Extended Data Table 2.** Total PNN counts in the hippocampus following exercise training.

| Total PNN count (PNN/mm<sup>2</sup>) |  |  |  |
| --- | --- | --- | --- |
|  | Sedentary (SD) | Exercise (SD) | Independent t test |
| CA1 | 0.343 (0.056) | 0.283 (0.061) | $t_{(4)} = 1.238, p = 0.283$ |
| CA2 | 0.163 (0.025) | 0.160 (0.022) | $t_{(4)} = 0.164, p = 0.878$ |
| CA3 | 0.221 (0.062) | 0.226 (0.037) | $t_{(4)} = -0.099, p = 0.926$ |
| CA3 (Pyramidal) | 0.256 (0.029) | 0.227 (0.096) | $t_{(4)} = 0.505, p = 0.640$ |
| CA4 | 0.573 (0.060) | 0.578 (0.032) | $t_{(4)} = -0.133, p = 0.900$ |
| DG | 0.247 (0.054) | 0.207 (0.038) | $t_{(4)} = 1.043, p = 0.356$ |

PNN count values, standard deviation, and statistical analyses reported with 3 decimal places. PNN – Perineuronal net. DG – dentate gyrus. $n = 3$ .

