## Supplementary material for "Exercise remodels hippocampal extracellular matrix to alleviate chondroitin-4-sulphate–induced memory impairment"

### Supplementary methods (submit as one PDF)

#### Supplementary Table 1. Treadmill familiarisation protocol.

| Minutes | Day 1 | Day 2 | Day 3 | Day 4 | Day 5 |
|---|---|---|---|---|---|
| 0-1 | 0 cm.s$^{-1}$ | 20 cm.s$^{-1}$ | 25 cm.s$^{-1}$ | 25 cm.s$^{-1}$ | 25 cm.s$^{-1}$ |
| 1-2 | 0 cm.s$^{-1}$ | 25 cm.s$^{-1}$ | 30 cm.s$^{-1}$ | 30 cm.s$^{-1}$ | 30 cm.s$^{-1}$ |
| 2-3 | 20 cm.s$^{-1}$ | 30 cm.s$^{-1}$ | 35 cm.s$^{-1}$ | 35 cm.s$^{-1}$ | 35 cm.s$^{-1}$ |
| 3-4 | 20 cm.s$^{-1}$ | 35 cm.s$^{-1}$ | 40 cm.s$^{-1}$ | 40 cm.s$^{-1}$ | 40 cm.s$^{-1}$ |
| 4-4:30 | 25 cm.s$^{-1}$ | 40 cm.s$^{-1}$ | 45 cm.s$^{-1}$ | 45 cm.s$^{-1}$ | 45 cm.s$^{-1}$ |
| 4:30-5 | 25 cm.s$^{-1}$ | 40 cm.s$^{-1}$ | 50 cm.s$^{-1}$ | 50 cm.s$^{-1}$ | 50 cm.s$^{-1}$ |

### Supplementary Table 2. TaqMan™ gene expression assay information (RhoA/ROCK pathway).

| Gene name | Gene type | TaqMan gene symbol | TaqMan assay ID | Amplicon length |
|---|---|---|---|---|
| 18s | Reference | 18s rRNA | Hs99999901_s1 | |
| Cyclophilin A (CyPA) | Reference | Ppia | Rn00690933_m1 | |
| Nogo-A | Target | Rtn4 | Rn00582903_m1 | 90 |
| Myelin-associated glycoprotein (MAG) | Target | Mag | Rn01457782_m1 | 56 |
| Oligodendrocyte-myelin glycoprotein (OMGp) | Target | Omg | Rn02533851_s1 | 131 |
| Semaphorin-4D (SEMA4D) | Target | Sema4d | Rn01435039_m1 | 62 |
| Ephrin-B3 (EFNB3) | Target | Efnb3 | Rn01750591_g1 | 82 |
| Aggrecan (ACAN) | Target | Acan | Rn00573424_m1 | 74 |
| Versican (VCAN) | Target | Vcan | Rn01493755_m1 | 89 |
| Neurocan (NCAN) | Target | Ncan | Rn00581331_m1 | 59 |
| Brevican (BCAN) | Target | Bcan | Rn00563814_m1 | 71 |
| Nogo receptor 1 (NgR1) | Target | Rtn4r | Rn00586061_s1 | 57 |
| Nogo receptor 2 (NgR2) | Target | Rtn4rl2 | Rn00710574_m1 | 63 |
| Nogo receptor 3 (NgR3) | Target | Rtn4rl1 | Rn01466695_m1 | 119 |
| Tumour necrosis factor receptor superfamily, member 19 (TROY) | Target | Tnfrsf19 | Rn01534699_m1 | 60 |
| Leucine rich repeat and Ig domain containing 1 (LINGO-1) | Target | Lingo1 | Rn03993618_s1 | 109 |
| Lysophosphatidic acid receptor 1 (LPAR1) | Target | Lpar1 | Rn00588435_m1 | 67 |
| Protein tyrosine phosphatase, receptor type, F (PTPRF) | Target | Ptprf | Rn00695914_m1 | 70 |
| Protein tyrosine phosphatase, receptor type, S (PTPRS) | Target | Ptprs | Rn00569511_m1 | 76 |
| Rho-associated coiled-coil containing protein kinase 2 (ROCK2) | Target | Rock2 | Rn00564633_m1 | 73 |
| Guanine nucleotide binding protein, alpha 13 (GNA13) | Target | Gna13 | Rn01461471_m1 | 106 |
| Rho guanine nucleotide exchange factor 12 (LARG) | Target | Arhgef12 | Rn01417838_m1 | 101 |
| LIM domain kinase 1 (LIMK1) | Target | Limk1 | Rn01499352_m1 | 69 |
| Myosin light chain 2 (MYL2) | Target | Myl2 | Rn01480558_g1 | 94 |
| Phosphatase and tensin homolog (PTEN) | Target | Pten | Rn00477208_m1 | 73 |
| Collapsin response mediator protein 2 (CRMP2) | Target | Dpysl2 | Rn01534654_m1 | 73 |
| Cofilin 1 (CFL1) | Target | Cfl1 | Rn01501422_g1 | 77 |

### Supplementary Table 3. Gene expression group numbers after data exclusions.

| Target gene | Hippocampus | | Cortex | | Lumbar | |
|---|---|---|---|---|---|---|
| | Sedentary (*n*) | Exercise (*n*) | Sedentary (*n*) | Exercise (*n*) | Sedentary (*n*) | Exercise (*n*) |
| ACAN | 3 | 3 | 3 | 2 | 4 | 4 |
| BCAN | 4 | 4 | 4 | 3 | 4 | 4 |
| CFL1 | 4 | 4 | 4 | 3 | 4 | 4 |
| DPYSL2 | 4 | 4 | 4 | 3 | 4 | 4 |
| EFNB3 | 4 | 4 | - | - | - | - |
| GNA13 | 4 | 4 | 4 | 3 | 4 | 4 |
| LARG (ARHGEF12) | 4 | 4 | 4 | 3 | 4 | 4 |
| LIMK1 | 2 | 2 | 4 | 3 | 4 | 4 |
| LINGO1 | 4 | 4 | 4 | 3 | 4 | 4 |
| LPAR1 | 4 | 4 | 4 | 3 | 4 | 4 |
| MAG | 4 | 4 | 4 | 3 | 4 | 4 |
| MYL2 | - | - | - | - | 3 | 4 |
| NCAN | 4 | 4 | 4 | 3 | 4 | 4 |
| NgR1 (RTN4R) | 4 | 4 | 4 | 3 | 4 | 4 |
| NgR2 (RTN4RL3) | 4 | 2 | 3 | 3 | - | - |
| NgR3 (RTN4RL1) | 4 | 3 | 3 | 3 | - | - |
| NOGO-A (RTN4) | 4 | 4 | 4 | 3 | 4 | 4 |
| OMG | 4 | 4 | 4 | 3 | 4 | 4 |
| PTEN | 4 | 4 | 4 | 3 | 4 | 4 |
| PTPRF | 2 | 2 | 2 | 2 | 3 | 3 |
| PTPRS | 3 | - | 3 | 3 | - | - |
| ROCK2 | 4 | 4 | 4 | 3 | 4 | 4 |
| SEMA4D | 4 | 4 | 4 | 3 | 4 | 4 |
| TROY (TNFRSF19) | 4 | 4 | 4 | 3 | 4 | 4 |
| VCAN | 4 | 4 | 4 | 3 | 4 | 4 |

Gene expression values were excluded if outside two standard deviations from the mean or if the Cq value was above cycle 35.

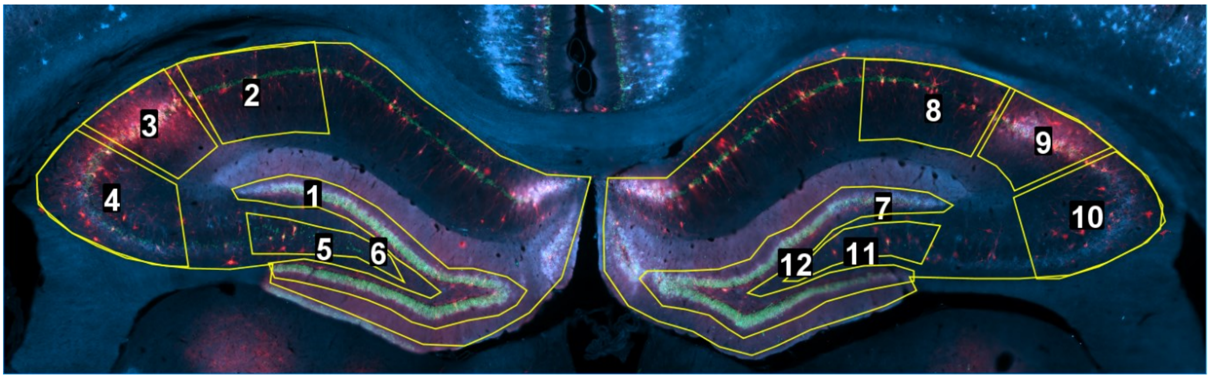


**Supplementary Figure 1. Hippocampal regions of interest.**

Hippocampal regions of interest manually drawn using ImageJ. 1 – Left total hippocampus (the number label is in the middle of the whole outlined region); 2 – Left CA1; 3 – Left CA2; 4 – Left CA3; 5 – Left CA4; 6 – Left dentate gyrus; 7 - Right total hippocampus (the number label is in the middle of the whole outlined region); 8 – Right CA1; 9 – Right CA2; 10 – Right CA3; 11 – Right CA4; 12 – Right dentate gyrus. Regions outlined using Figure 33 (Bregma -3.30 mm) in the Rat Brain Atlas (Paxinos and Watson, 2013).

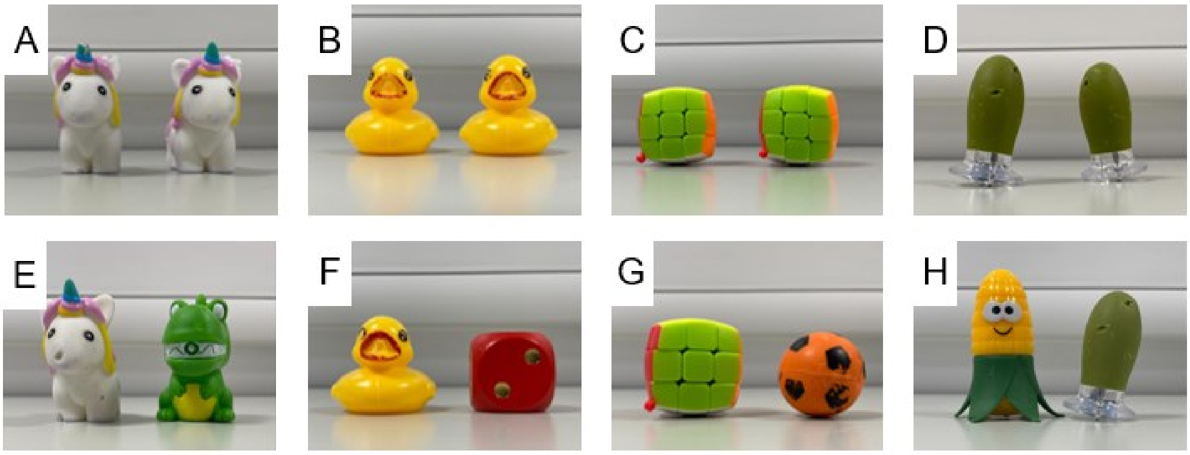


**Supplementary Figure 2. Object pairs used for novel object recognition tests.**

A-D. Identical object pairings used in the five-minute sample phase. E-H. Novel object pairings used in the three-minute choice phase.
